# MFSD7c functions as a transporter of choline at the blood-brain barrier

**DOI:** 10.1101/2023.10.03.560597

**Authors:** Xuan T.A. Nguyen, Thanh Nha Uyen Le, Hoa Thi Thuy Ha, Toan Q. Nguyen, Anna Artati, Nancy C.P. Leong, Dat T. Nguyen, Pei Yen Lim, Adelia Vicanatalita Susanto, Huang Qianhui, Fam Ling, Leong Lo Ngah, Isabelle Bonne, Angela Lee, Catherine Gooch, Jorge Luis Granadillo De Luque, Dejie Yu, Huang Hua, Tuck Wah Soong, Matthew Wook Chang, Markus R. Wenk, Jerzy Adamski, Amaury Cazenave-Gassiot, Long N. Nguyen

## Abstract

Mutations of MFSD7c (also known as *Flvcr2*), which is an orphan transporter, are linked to Fowler syndrome ^1, 2^. Here, we use *Mfsd7c* knockout mice and cell-based assays to reveal that MFSD7c is a choline transporter at the blood-brain barrier (BBB). We performed comprehensive metabolomics and detected differential changes of metabolites in the brains and livers of *Mfsd7c* knockout (*Mfsd7c*^−/−^) embryos. Particularly, we found that choline-related metabolites were altered in the brains but not in the livers of *Mfsd7c*^−/−^ embryos. Thus, we hypothesized that MFSD7c regulates the levels of choline in the brain. Indeed, expression of human *MFSD7c* in cells significantly increased choline uptake. Interestingly, we showed that choline uptake by *MFSD7c* is greatly increased by choline-metabolizing enzymes, leading us to demonstrate that MFSD7c is a facilitative transporter of choline. Furthermore, single-cell patch-clamp showed that the import of choline by MFSD7c is electrogenic. Choline transport function of MFSD7c is conserved in vertebrates, but not in yeasts. We show that human MFSD7c is a functional ortholog of HNM1, the yeast choline importer. Employing our transport assays, we showed that several missense mutations of human *MFSD7c* from Fowler patients had abolished or reduced choline transport activity. Mice lacking *Mfsd7c* in the CNS endothelial cells suppressed the import of exogenous choline from blood but unexpectedly had increased choline levels in the brain. Stable-isotope tracing study revealed that MFSD7c is required for exporting choline derived from lysophosphatidylcholine (LPC) in the brain. Collectively, our work identifies MFSD7c as a choline transporter at the BBB. This study suggests that defective export of choline in the brain may be a cause of Fowler syndrome.

## Introduction

Missense mutations of *MFSD7c* (also known as *FLVCR2*) have been described in Fowler patients ^2^. The major clinical symptoms reported in these patients are the severe dilation of cerebral blood vessels and mild microcephaly. Most severe cases result in embryonic death, but some patients have survived after birth ^1, 2, 3, 4, 5, 6^. Although the cause of Fowler syndrome is unclear, it is linked to the loss of functions of MFSD7c, an orphan transporter. To gain better understanding about the disease mechanisms, the molecular role of MFSD7c must be elucidated. We recently reported that mice with a global deletion of *Mfsd7c* had late gestation lethality with severe dilation of central nervous system (CNS) vasculature ^1^. Specific deletion of *Mfsd7c* in endothelial cells also resulted in similar phenotypes, indicative of its critical roles in the blood vessels ^7^. *Mfsd7c* knockout embryos exhibited multiple neurological phenotypes that are probably linked to primary defects of the CNS vasculature. These phenotypes resemble the glomeruloid structures in the CNS blood vessels of patients with Fowler syndrome ^1, 7^.

MFSD7c belongs to the Major Facilitator Superfamily (MFS) of transporters which facilitate the movement of small molecules through cell membranes ^8^. MFSD7c was reported to exert heme import activity ^9^, but this observation has not been confirmed. Recently, it was shown that MFSD7c is involved in thermogenesis in response to heme. Besides its localization in the mitochondria, this study and our results also showed that MFSD7c is expressed in the plasma membrane ^1, 10^. Furthermore, we and others showed that MFSD7c is expressed in endothelial cells of the CNS vasculature ^1, 7^. These results suggest that MFSD7c is a membrane transporter for small molecules at the blood-brain barrier (BBB). Utilizing mouse knockout models coupled with comprehensive metabolomics, we revealed the specific changes in metabolites due to the loss of *Mfsd7c*. These results guide us to demonstrate that MFSD7c is a transporter for choline. We show that MFSD7c facilitates choline transport via the plasma membrane in a concentration-dependent manner. Unexpectedly, we find that lack of Mfsd7c at the BBB causes choline accumulation in the brain. Stable isotopic tracing study reveals that lysophosphatidylcholine (LPC) and Mfsd2a pathway delivers choline to the brain. The excessive amount of choline is then released to be exported out of the brain via MFSD7c. The current work identifies MFSD7c as a choline transporter and provides a foundation that facilitates the understanding of the disease mechanisms of Fowler syndrome. Additionally, the identification of MFSD7c as a choline transporter at the BBB lays the groundwork for future research to reveal the metabolic fates of choline in the brain.

## Results

### Lack of *Mfsd7c* affects choline levels in the brain

Deletion of *Mfsd7c* resulted in late gestation lethality in mice ^1, 7^. Because MFSD7c is predicted to be a membrane transporter, its deletion is anticipated to cause insufficient transport of essential nutrients to the brain. To deorphanize the physiological ligands for MFSD7c, we performed a comprehensive analysis of metabolites in the brains of WT and global *Mfsd7c* knockout (*Mfsd7c*^−/−^) embryos at gestation day 14.5 (E14.5), a time point when knockout embryos already exhibited vascular growth defects^1^. This analysis encompassed more than 520 metabolites from metabolic pathways for carbohydrates, lipids, amino acids, and cofactors (**Supplemental data tables 1-10**). First, we found that the levels of glycolytic metabolites such as lactate were significantly increased, whereas the levels of 3-phosphoglycerate (3-PGA) and phosphoenolpyruvate (PEP) were reduced (**Figure 1a**). These metabolic changes are congruent with the increased glycolysis and possibly reduced mitochondrial activity as a consequence of hypoxia that occurs in the brain of *Mfsd7c*^−/−^ embryos^1,7^. MFSD7c was shown to regulate heme transport^10^. However, heme and bilirubin levels in brains were comparable between *Mfsd7c*^−/−^ and WT embryos (**Extended Data Figure 1a-b**). Second, we detected the increased levels of several long chain acylcarnitines (**Figure 1b, c**; black arrowheads), whereas the levels of L-carnitine were significantly reduced in the brain of *Mfsd7c*^−/−^ embryos (**Figure 1d**). Nevertheless, we ruled out that MFSD7c transports these acylcarnitines by performing the import assay with palmitoylcarnitine (C16:0 carnitine) and octanoylcarnitine (C8:0 carnitine) (**Extended Data Figure 1c-d**). Again, these changes might reflect the defects of mitochondrial activity due to hypoxia in the knockout brain. Indeed, mitochondrial morphology and the expression level of several mitochondrial markers was reduced in the brain of *Mfsd7c*^−/−^ embryos (**Extended Data Figure 2**). Interestingly, we noted the increased levels of choline, palmitoylcholine, and oleoylcholine in the brains of *Mfsd7c*^−/−^ embryos relative to WT embryos (**Figure 1b; red arrowheads**). In contrast, the levels of CDP-choline were reduced in *Mfsd7c*^−/−^ embryos (**Figure 1d**). These changes in choline metabolites were specific to the brain as their levels were unaltered in the livers from *Mfsd7c*^−/−^ embryos (**Figure 1e**). Instead, we detected a striking increase in the levels of several neutral lipids such as monoacylglycerols and diacylglycerols in the livers of *Mfsd7c*^−/−^ embryos (**Figure 1f, arrowheads**). Deficiency of choline has been linked to mitochondrial defects ^11, 12, 13, 14^. These differential changes in the levels of choline and choline metabolites in the brain and the accumulation of neutral lipids in the livers of *Mfsd7c*^−/−^ embryos suggest that MFSD7c may be involved in choline metabolism.

**Figure 1.**
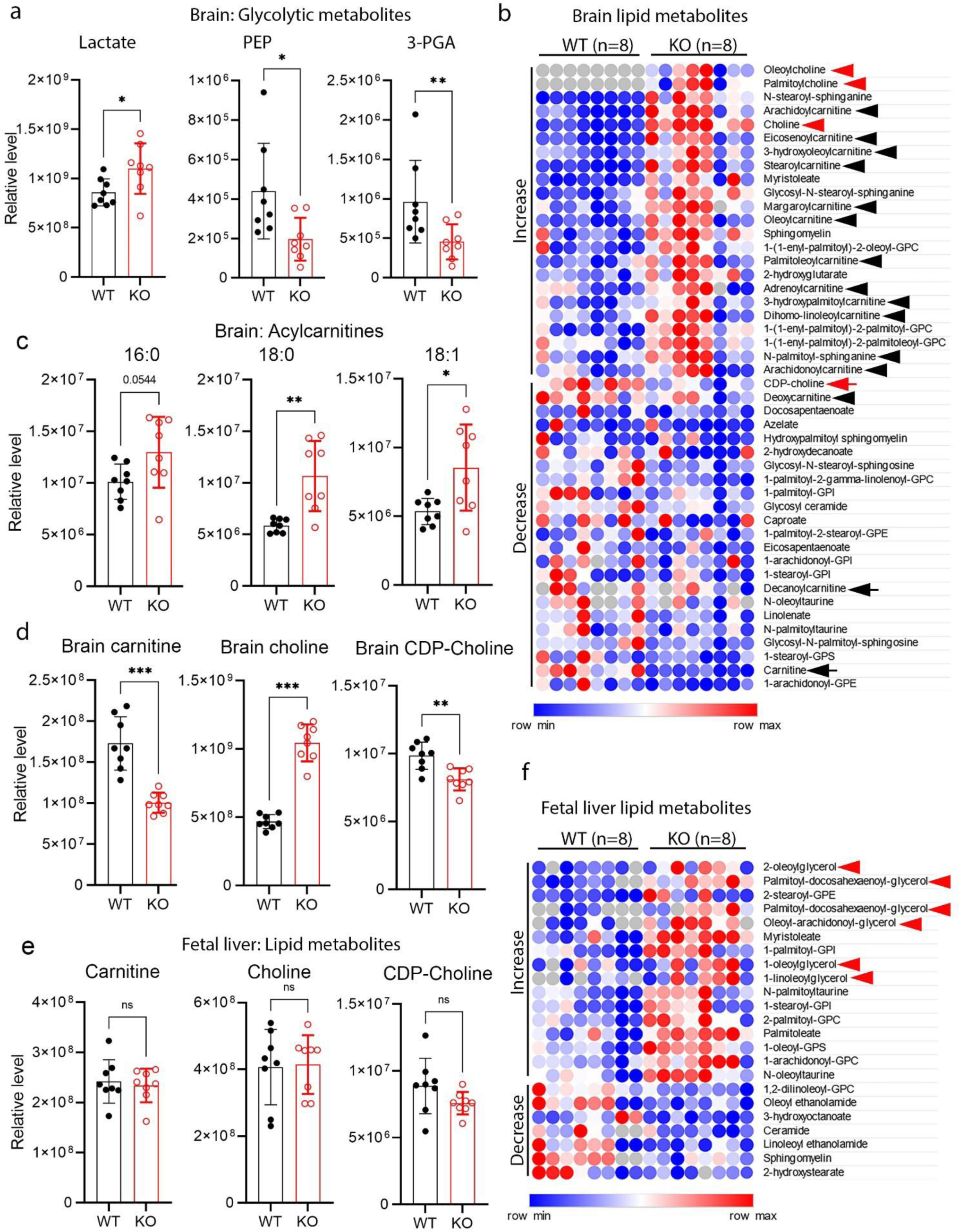
Comprehensive metabolomics identifies changes in the levels of choline and choline metabolites in the brain of *Mfsd7c* knockout. **a**, Increased levels of glycolytic metabolite lactate and decreased levels of mitochondrial metabolites in Mfsd7c knockout brain. **b,** Heatmap of metabolites with differential changes in the brains of Mfsd7c knockout compared to that of WT embryos. **c,** Increased levels of acylcarnitines in the brains of Mfsd7c knockout embryos. **d,** Increased levels of choline and decreased levels of CDP-choline and L-carnitine in the brains of Mfsd7c knockout embryos. **e,** Levels of L-carnitine, choline, and CDP-choline were unchanged in the fetal livers of WT and Mfsd7c knockout embryos. **f,** Heatmap of neutral lipid species that are differentially changed in the fetal livers of Mfsd7c knockouts compared to that of WT embryos. The results showed the accumulation of neutral lipids in the fetal livers of the Mfsd7c knockout embryos. The brains and livers were collected at E14.5. N=8 per genotype. ***P<0.001, **P<0.01, *P<0.05; non-parametric t-test (Mann Whitney test); ns, not significant. Data are mean and SD; each dot represents one mouse.

### MFSD7c mediates transport of choline

The accumulation of choline and decrease in CDP-choline levels in the brains of *Mfsd7c*^−/−^ embryos suggests that MFSD7c is important for regulation of choline levels. Thus, we tested whether MFSD7c regulates choline levels by a transport mechanism. We overexpressed human (*hMFSD7c*) or mouse *Mfsd7c* (*mMfsd7c*) in HEK293 cells and utilized radioactive [^3^H]-choline for transport assays. Interestingly, overexpression of human or mouse MFSD7c led to 1.5-2 fold increase of intracellular levels of radioactive signals, implicating that MFSD7c mediates the import of choline (**Figure 2a**). We validated our choline transport assay by including neuronal choline transporter CHT1 (also known as SLC5a7) as a positive control (**Figure 2a**). The import of choline mediated by hMFSD7c was increased with the increased levels of choline in medium and was slightly increased over time (**Figure 2b-c**). To reaffirm that MFSD7c mediates choline import to the cells, we included the missense mutation, S203Y in our transport assays. In comparison to wild-type MFSD7c, S203Y mutant exhibited abolished choline transport activity when incubated with different concentrations of choline (**Figure 2b-c**). The reduced import of choline of the wildtype and mutant was not due to reduced protein expression levels nor defective localization in the plasma membrane (**Extended Data Figure 3a-b**). MFSD7c is conserved from fish to man, but not in yeasts (**Extended Data Figure 3c**). We tested choline import function for several *MFSD7c* orthologs including the ones from zebrafish (*DaMfsd7c_a* and *DaMfsd7c_b* isoforms), medaka fish (*MeMfsd7c*), and frog (*XeMfsd7c)*. Consistently, these MFSD7c orthologs exhibited equal choline transport activity compared to hMFSD7c (**Figure 2d**). In the yeast *S. cerevisiae*, HNM1 was shown to import choline as well as L-carnitine^15^. We performed complementation assays of *hMFSD7c* in *Hnm1* S. *cerevisiae* knockout. First, we showed that *Hnm1* knockout cells exhibited reduced choline uptake compared to wild-type yeast cells (**Figure 2e**). Second, when wild-type yeast cells and *Hnm1* mutant yeast cells were overexpressed with human MFSD7c, they both exhibited significantly increased choline uptake compared to parental cells (**Figure 2e**, three different clones were used**; also Extended Data Figure 4a-b**). These results indicate that *hMFSD7c* can rescue the loss of yeast *Hnm1* and reaffirm that MFSD7c is a choline transporter. Choline transport by neuronal choline transporter CHT1 is inhibited by hemicholinium-3 (HC3), a competitive inhibitor of CHT1^16,17^. Hemicholinium-3 inhibited CHT1, but not MFSD7c, suggesting that these two choline transporters exert different transport properties (**Figure 2f**). SLC44a1 and SLC44a2 has been reported to be choline transporters. However, we were unable to show choline import activity for these transporters under the tested conditions (**Extended Data Figure 4c-d**). MFSD7c prefers choline as a substrate as it did not transport betaine, a choline metabolite that is also found in the brain (**Figure 2g-h**). Similarly, *Mfsd7c* did not transport L-carnitine, acetyl-carnitine, and ethanolamine, and acetylcholine (**Figure 2i-k; Extended Data Figure 4e**). Together, our results reveal that MFSD7c is a conserved transporter for choline in vertebrates.

**Figure 2.**
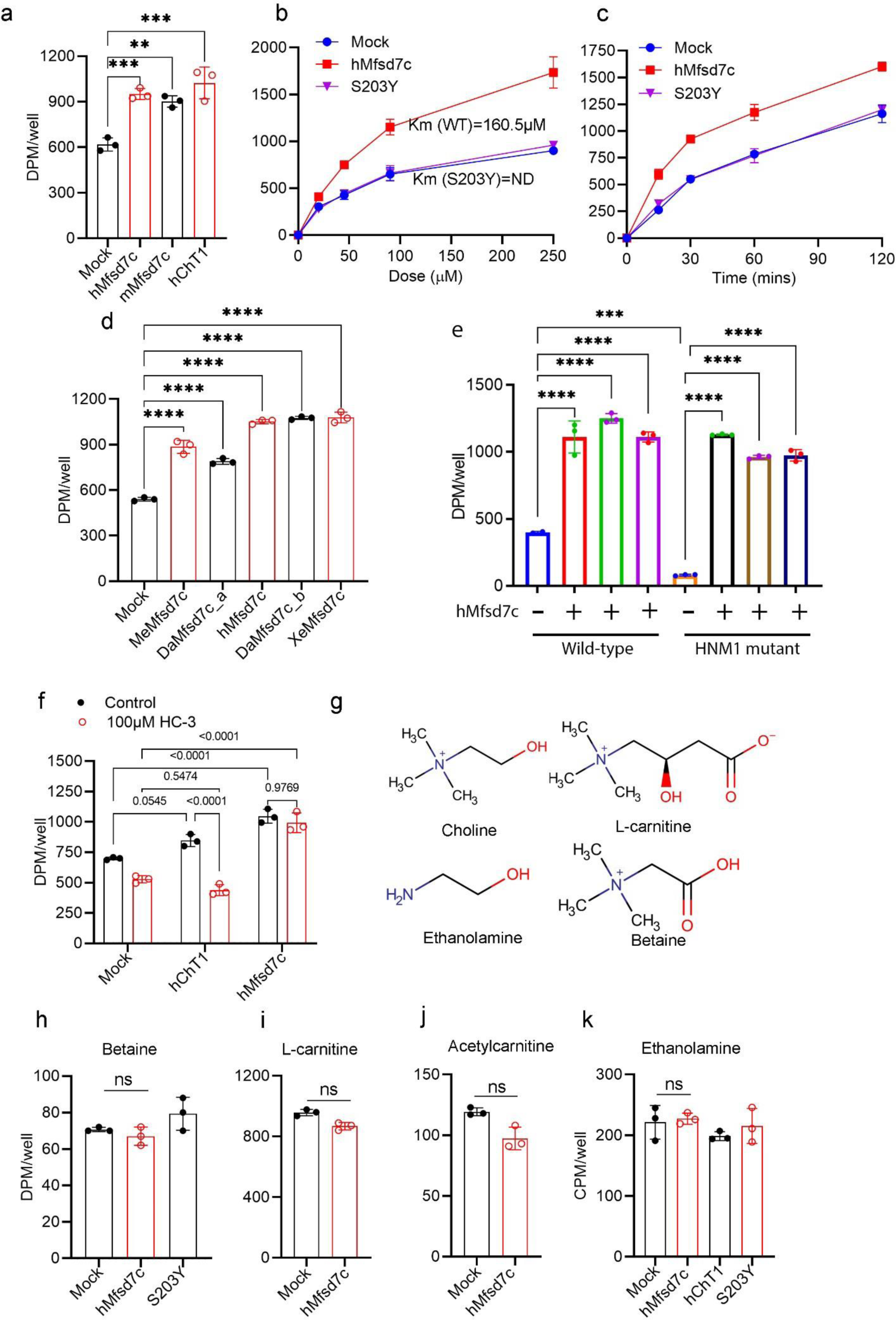
Expression of MFSD7c increases choline uptake. **a,** Overexpression of mouse Mfsd7c (mMfsd7c) and human Mfsd7c (hMfsd7c) increases choline uptake in HEK293 cells. CHT1 (SLC5a7), a neuronal choline transporter, was included as a positive control. **b**, Dose curve for choline uptake from mock, hMfsd7c, and S203Y mutant. **c**, Time course for choline uptake from mock, hMfsd7c, and S203Y mutant. Experiments were performed at least twice in triplicates. **d**, Transport activity of mock, medaka Mfsd7c (MeMfsd7c), zebrafish Mfsd7c isoform a (DaMfsd7c_a), hMfsd7c, zebrafish Mfsd7c isoform b (DaMfsd7c_b), and frog Mfsd7c (XeMfsd7c). **e**, Complementation assays of hMfsd7c with the knockout of yeast S. cerevisiae choline transporter HNM1. Transport activity of wild-type S. cerevisiae strain and HNM1 mutant strain with and without overexpression of hMfsd7c. The results show that expression of hMfsd7c overcomes the loss of choline transport activity in HNM1 mutant yeast cells. Three different overexpression yeast clones were used. **f**, Choline transport activity by CHT1 and hMfsd7c in the presence of hemicholinium-3 (HC-3), a competitive inhibitor of CHT1. **g**, Structure of choline, betaine, L-carnitine, and ethanolamine. **h-k**, Uptake assays of hMfsd7c, S203Y, and/or CHT1 with betaine, L-carnitine and acetylcarnitine, and ethanolamine. Each symbol represents one biological replicate. All of these experiments were repeated at least twice in triplicate. Data are expressed as mean ± SD. ****P<0.0001, ***P<0.001, **P<0.01, ns: not significant. One-way ANOVA for a, d, e, f, h, and k; Two-way ANOVA for b and c; t-test for i and j.

### MFSD7c functions as a facilitative transporter for choline

We noted that the transport activity of MFSD7c was slightly enhanced over time (**Figure 2c**), suggesting that intracellular levels of choline might affect the influx of choline. Thus, we characterized its transport properties by testing whether MFSD7c utilizes cations for transport activity. Replacement of sodium with lithium did not affect choline import by MFSD7c (**Figure 3a; Extended Data Figure 5a**). Next, we tested whether lowering intracellular choline levels could increase the influx of choline via MFSD7c. To test this hypothesis, we co-expressed *hMfsd7c* with human choline kinase A (hChKA), which phosphorylates choline to phosphorylcholine, thus lowering the intracellular concentration of choline. Remarkably, co-expression of *hMfsd7c* with hChKA increased choline uptake by approximately 9-fold (**Figure 3b**), whereas sole expression of *hMfsd7c* increased choline uptake by 1.5-2 fold (**Figure 2a**). In the presence of hChKA, the import of choline was significantly increased over time (**Figure 3c**). Co-expression of hChKA with *hMfsd7c* also greatly enhanced choline uptake in a dose-dependent manner (**Figure 3d; Extended Data Figure 5b**). The kinetics (Km) and Vmax for choline import by human Mfsd7c under this condition was 100µM and 0.117 µmol/well/60 mins, respectively. Choline uptake by S203Y mutant in the absence of hChKA was completely inhibited (**Figure 2b-c**). Interestingly, we noted that there was a slight increase in choline import in S203Y mutant when co-expressed with hChKA, indicating that the transport activity of S203Y is severely delayed, but not abolished (**Figure 3d; Extended Data Figure 5c**). The kinetics (Km) and Vmax for importing choline by S203Y mutant under this condition was 1197µM and 0.039 µmol/well/60 mins, respectively. We also examined whether expression of choline acetyltransferase (CHAT), a neuronal enzyme for acetylcholine synthesis, would increase choline uptake by converting choline into acetylcholine. Indeed, co-expression of CHAT and *hMfsd7c* greatly increased choline uptake (**Figure 3e**). In the absence of ETNK1, an ethanolamine kinase, MFSD7c did not increase uptake of ethanolamine (above **Figure 2k**). However, transport of ethanolamine by MFSD7c was increased to a significant level when ETNK1 was co-expressed (**Figure 3f**). Interestingly, co-expression of *hMfsd7c* with hChKA slightly increased L-carnitine uptake (**Figure 3g**). However, L-carnitine is unable to compete for choline import, suggesting that L-carnitine is a weak ligand (**Extended Data Figure 5d**). Furthermore, we tested whether the transport of choline by MFSD7c is bidirectional. Indeed, our results showed that expression of *hMfsd7c* is required for the release of intracellular choline (**Figure 3h**). Choline is a positively charged molecule. Import of choline would increase the membrane potential. To this end, we performed whole-cell patch clamp of HEK293 cells co-expressing *hMfsd7c* or S203Y mutant with hChKA. Cells with overexpression of hChKA alone were used as a control (**Extended Data Figure 5e**). Incubation of control cells or cells expressing S203Y mutant with 100µM or 200µM choline did not significantly increase membrane potential (**Figure 3i**). In contrast, expression of *hMfsd7c* significantly increased membrane potential when 100µM or 200µM choline was included in the bath solution (**Figure 3i, j**). Interestingly, removal of choline reversed the increased membrane potential, suggesting that the influx of choline generates the membrane potential. These results show that the import of choline by MFSD7c is electrogenic. Collectively, our results show that MFSD7c facilitates the transport of choline and that it behaves like a channel for the movement of choline via the plasma membrane.

**Figure 3.**
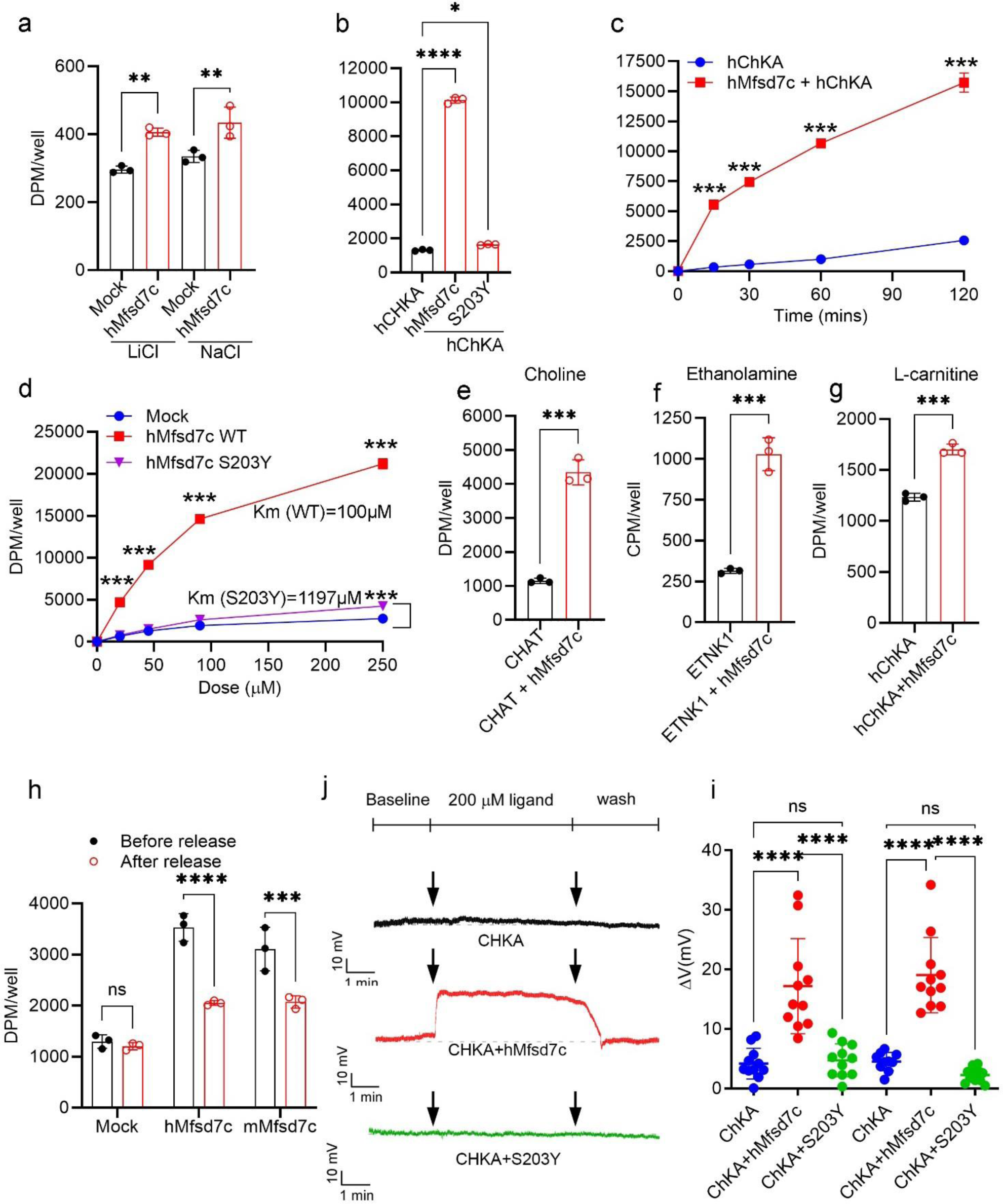
MFSD7c exhibits properties of a facilitative transporter for choline. **a,** Transport activity of hMfsd7c is not sodium-dependent. **b**, Import of choline by hMfsd7c is strongly increased by co-expression of human choline kinase A (hChKA). **c**, Time course of choline uptake by hMfsd7c when co-expressed with hChKA. **d**, Dose curve of choline uptake by hMfsd7c and S203Y mutant when co-expressed with hChKA. Data are expressed as mean ± SD. These experiments were repeated at least twice in triplicate. ***P< 0.0001, **P< 0.01, *P< 0.05. One-way ANOVA: a and b. Two-way ANOVA: c and d. **e**, Import of choline by hMfsd7c is also significantly increased with co-expression of choline acyltransferase (CHAT). **f**, Co-expression of hMfsd7c with ethanolamine kinase 1 (ETNK1) increases uptake of ethanolamine. **g**, Co-expression of hMfsd7c with hChKA slightly increased uptake of L-carnitine. **h**, hMfsd7c or mMfsd7c mediates the release of choline from cells. Before release: the levels of choline in the cells; After release: the remaining levels of choline in the cells after 30 mins incubation with plain DMEM. There was a reduction of intracellular choline due to the release into the medium. Data are expressed as mean ± SD. These experiments were repeated at least twice in triplicate. ****P< 0.0001, ***P< 0.001, t-test: e-g; One-way ANOVA: h. **i**, Uptake of choline by hMfsd7c increases membrane potential in HEK293 cells. Representative exemplar traces of HEK293 cells co-transfected with empty pIRES2-eGFP plasmid and hChKA plasmid (top), hChKA with hMfsd7c (middle), and hChKA with S203Y (bottom). The effect of 200 µM choline on the resting membrane potential changes was induced by active import of choline by hMfsd7c, but not mutant S203Y. **j**, Quantification of resting membrane potential changes induced with 100 and 200 µM choline. Data are expressed as mean ± SD. ****P< 0.0001; ns, not significant; One-way ANOVA.

### Lack of Mfsd7c results in elevated levels of choline in the brain

Choline uptake at the BBB has been documented in previous studies^18,19,20^. However, the identity of the long-sought choline transporter was unknown. To test if MFSD7c play a role for choline transport *in vivo*, we generated conditional knockout mice for *Mfsd7c* in the endothelial cells using Cdh5-CreERT^2^ (**Figure 4a**). Next, we intravenously injected controls (*Mfsd7c*^f/f^) and *Mfsd7c*^f/f^Cdh5-CreERT^2^ (hereafter, Ec*Mfsd7c*-KO) mice with a dose (2mM in blood) of radioactive choline and measured the distribution of radioactive signals in the brains, lungs, livers, hearts, and kidneys after 30 minutes. We chose this supraphysiological dose as the half-life of choline in plasma is within 3 minutes^21^. The radioactive choline levels in blood were indifferent between the two genotypes after 5 mins choline injection, indicating that choline was equally delivered to the blood of the mice (**Figure 4b**). Importantly, we found that radioactive signals were significantly reduced in the brains of Ec*Mfsd7c*-KO mice compared to control mice (**Figure 4c**). However, we noted that the majority of injected choline was present in peripheral organs such as livers and kidneys; and the radioactivity in the organs was comparable between the genotypes, indicating that reduced choline import to the brain was specific in Ec*Mfsd7c*-KO mice (**Figure 4d-g**). These results indicate that *Mfsd7c* can mediate the uptake of exogenous choline at the blood-brain barrier, a result which is consistent with the reported findings in the literature^18,19,20^. Nevertheless, the injected amount of choline was well above physiological concentrations of choline in blood ^22,23^. Perhaps, Mfsd7c mediates to import a small amount of choline to the brain at this supraphysiological dose.

**Figure 4.**
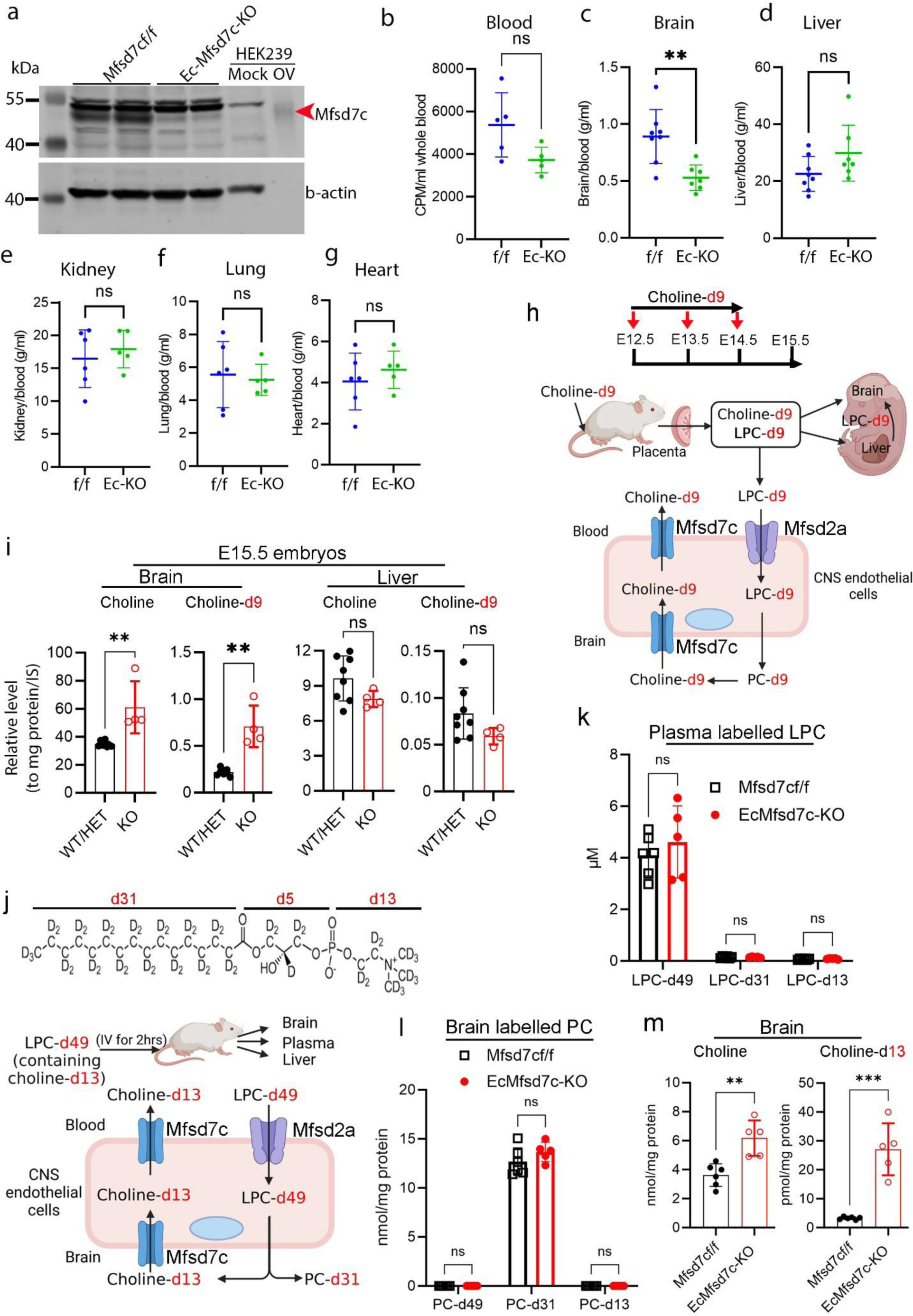
MFSD7c regulates choline levels in the brain. **a**, Western blot analysis of the micro-vessels in the brains. Mfsd7c band was reduced in EcMfsd7c-KO compared to controls (Mfsd7c^f/f^). Mock: 15µg proteins from mock transfected HEK239 cells; OV: 1.5µg protein lysates from overexpression of hMfsd7c in HEK293 cells used for interpretation of Mfsd7c band. **b**, Radioactive signals in the blood at 5 mins post-injection of radioactive choline in EcMfsd7c-KO (Ec-KO) and control (f/f) mice**. c-g**, The radioactive signals in the brain (**c**), livers (**d**), kidneys (**e**), lungs (**f**), and hearts (**g**) of EcMfsd7c-KO and control mice after 30 mins of choline injection. There was a significant reduction of radioactive signals in the brain of EcMfsd7c-KO mice compared to controls. Each symbol represents one biological replicate. **P< 0.01; t-test. **h**, Illustration of maternal deuterated choline delivery to the embryos. Choline-d9 is used for the synthesis of LPC-d9 by the maternal liver which is then delivered to the embryos. **i**, Levels of endogenous choline and choline-d9 in the brains and livers of WT/HET and KO embryos. Endogenous choline and choline-d9 were significantly increased in the brains but not in the liver of KO embryos. Each symbol represents one mouse. **P< 0.01; *P< 0.05, t-test. **j**, Illustration of deuterated LPC-d49 tracing experiment in adult mice. Plasma LPC-d49 is taken up to the brain by Mfsd2a for PC synthesis. Choline-d13 released in the brains after the remodelling of LPC/PC-d49 is imported back to the endothelial cells by Mfsd7c. **k,** Levels of LPC-d49, LPC-d31, and LPC-d13 the plasma of controls and EcMfsd7c-KO mice at 2hr post-injection of LPC-d49**. l,** Levels of PC-d49, PC-d31, and PC-d13 in the brains of controls and EcMfsd7c-KO mice at 2hr post-injection of LPC-d49. **m,** Levels of endogenous choline and choline-d13 in the brains of controls and EcMfsd7c-KO mice at 2hr post-injection of LPC-d49. Each symbol represents one mouse. \*\**P< 0.01,* \*\*\**P< 0.001; t-test and two-way ANOVA (in k and l)*.

Our metabolomic analysis showed that the levels of choline in the brain of *Mfsd7c^−/−^* embryos were elevated (**Figure 1d**). Mfsd7c is expressed in both sides of the CNS endothelial cells, it might suggest that Mfsd7c is also required for choline import from the brain parenchyma ^1^. To gain insights into the mechanism by which choline level is elevated in the brain of *Mfsd7c^−/−^* embryos, we injected deuterated choline (choline-d9) to the circulation of *Mfsd7c* pregnant dams and harvested *Mfsd7c^−/−^*embryos for lipidomic analysis of choline-containing lipids and choline (**Figure 4h**). We found that the total levels of endogenous phospholipids such as lysophosphatidylcholine (LPC), phosphatidylcholine (PC), and sphingomyelin (SM) as well as the deuterated phospholipids containing choline-d9 the brains and livers of *Mfsd7c* knockout and control littermates were comparable (**Supplemental Data table 11**). Interestingly, we found that the levels of endogenous choline and choline-d9 were significantly elevated in the brains but not livers of *Mfsd7c* KO embryos (**Figure 4i; Supplemental Data table 11**). Free choline import to the brain is expected to be suppressed in the KO embryos. These results suggest that choline imported to the brains of *Mfsd7c* KO embryos must be via other routes. We suspect that lysophosphatidylcholine (LPC) delivers choline to the brain of *Mfsd7c^−/−^* embryos (**Figure 4h**). These above results suggest a possibility that MFSD7c is also necessary for the export of choline from the brain parenchyma.

Lysophosphatidylcholine is one of the most abundant lipids in the plasma^24,25^ and is imported to the brain by Mfsd2a via the BBB for phospholipid synthesis^26,27^. To test our hypothesis that choline from LPC is accumulated in the brain of *Mfsd7c* knockout mice, we injected a dose (300µM in blood) of deuterated LPC-d49 which consists of choline-d13, palmitate-d31, and glycerol-d5 into the circulation of Ec*Mfsd7c*-KO and control mice and employed mass spectrometry for detection of choline-d13 and deuterated LPC and PC (**Figure 4j**). In the plasma, a part of LPC-d49 was remodelled to produce LPC-d31 and LPC-d13. However, LPC-d49 was still the major deuterated LPC in the blood of the mice at 2 hours post-injection (**Figure 4k**). In the brain, we were able to detect deuterated PC species, which were present with similar amounts in the phospholipid pool of Ec*Mfsd7c*-KO and control mice. These results indicate that plasma LPC-d49 was equally imported to the brains for PC synthesis (**Figure 4l; Supplemental Data table 12**). Remarkably, we found that most of the deuterated PC species in the brains of the mice contained palmitate-d31 (PC-d31), instead of the intact form of LPC-d49 (PC-d49) (**Figure 4l; Supplemental Data table 12**). This indicates that there was a dramatic remodelling of plasma derived LPC in the brain. As a result, the levels of choline-d13 in the brains of Ec*Mfsd7c*-KO mice were significantly increased compared to that of control mice (**Figure 4m**). Consistently, the levels of endogenous choline were confirmed to be elevated in Ec*Mfsd7c*-KO in the same analysis (**Figure 4m; Extended Data Figure 6a-b**), whereas the levels of choline-d13 in the plasma and livers of Ec*Mfsd7c*-KO and control mice were comparable, highlighting a specific elevation of choline in the brain of Ec*Mfsd7c*-KO mice (**Supplemental Data table 12**). Furthermore, the levels of acetylcholine were increased in the brains of Ec*Mfsd7c*-KO adult mice, especially from mice fed with choline deficient diet (CDD) (**Extended Data Figure 6a-b**). Since free choline import to the brain is likely suppressed or reduced in Ec*Mfsd7c*-KO mice (See **Figure 4c**), these results reveal an unexpected finding that excessive choline from LPC/PC metabolism is to be transported out of the brain and this mechanism is mediated by the activity of MFSD7c at the BBB.

### Defective choline export in the brain may be a causal factor and the missense mutations of *MFSD7c* in Fowler patients result in reduced choline transport activity

At this point, it is unclear whether deficiency of choline is a causal factor for the phenotypes of *Mfsd7c^−/−^* embryos or it is due to choline accumulation in the brain. If choline is utilized for phospholipid synthesis, it is expected that the brain phospholipid profile of *Mfsd7c* knockout mice must be changed. We performed lipidomic analysis of the brains from *Mfsd7c* knockouts and controls. However, there was no significant changes in the levels of major phospholipids containing choline such as LPC, PC, and SM due to Mfsd7c knockout embryos and mice (**Extended Data Figure 7a-b; Supplemental data tables 13-16**). Additionally, we compared the brain transcriptomes of *Mfsd7c^−/−^* embryos to that of wild-type embryos obtained from dietary choline deficient mice as we argued that deficiency of choline would recapitulate the deletion of *Mfsd7c* (**Extended Data Figure 8; Supplemental data table 17**). Nevertheless, we did not observe any significant change in gene expression from wild-type embryos obtained from dams fed with choline deficient diet compared to that of *Mfsd7c^−/−^* embryos. These results indicate that deficiency of choline is unlikely to be the cause for the phenotypes in the brain of *Mfsd7c^−/−^* embryos. Next, we tested whether depletion of choline from diets would improve the phenotypes in Mfsd7c^−/−^ embryos. We depleted choline from the diets for pregnant *Mfsd7c* mice and examined the CNS blood vessels of *Mfsd7c^−/−^* embryos. Interestingly, we found that depletion of choline significantly reduced the dilation of blood vessels in the cortices of E14.5 *Mfsd7c^−/−^* embryos (**Figure 5a-e**). Nonetheless, maternal choline deficiency did not prevent the migratory defects of the blood vessels to the ganglionic eminence (GE) regions of these *Mfsd7c^−/−^* embryos (**Figure 5d**), probably due to the dominant source of choline from LPC. Although the dietary intervention approach did not completely reverse the phenotypes of *Mfsd7c^−/−^*embryos, these results suggest that accumulation of choline might be detrimental to the developing blood vessels. Together, our results point to a critical role of MFSD7c for the regulation of choline levels in the brain.

**Figure 5.**
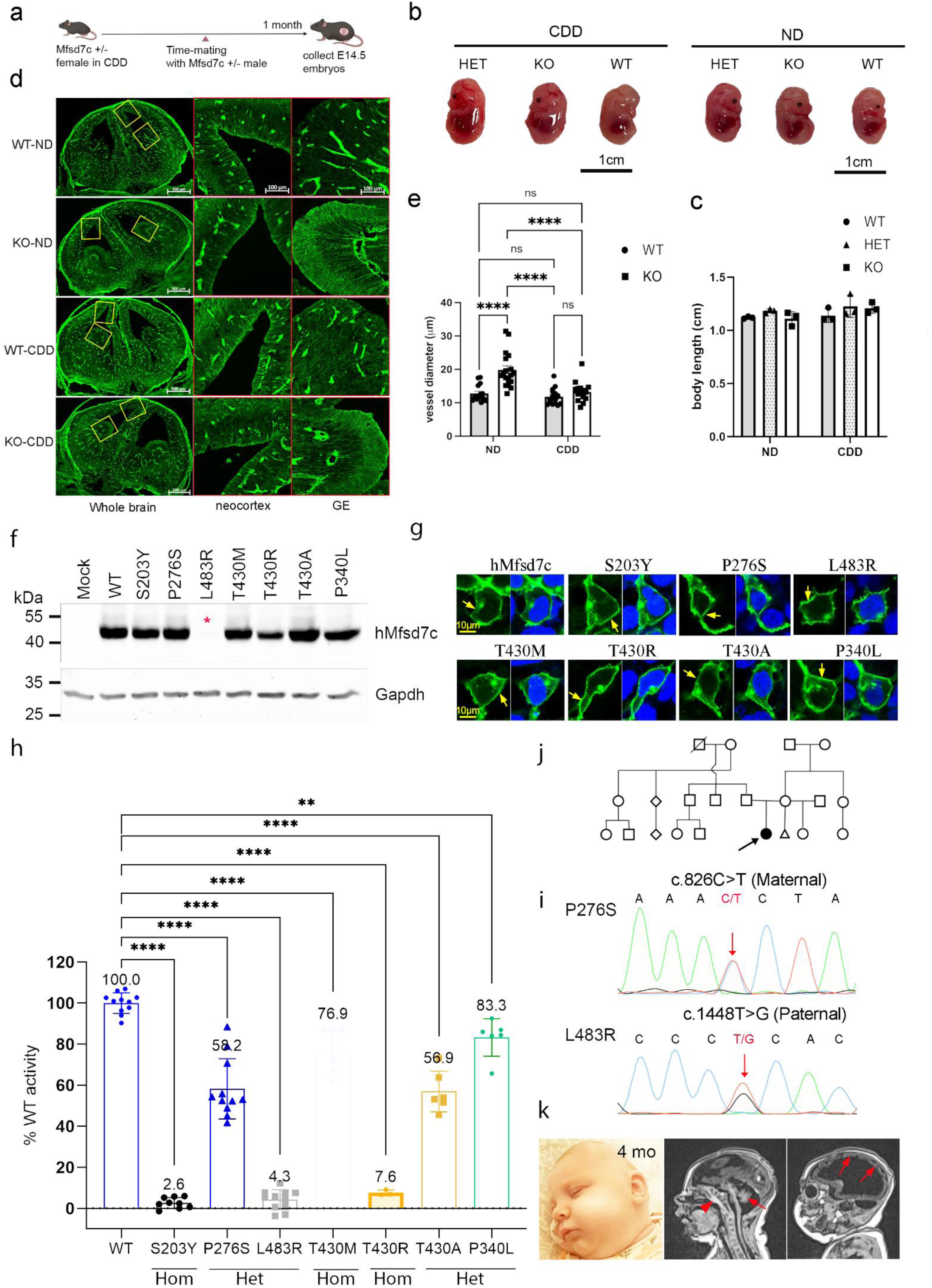
Defective choline export in the brain might confer to the phenotypes in the mice and the pathogenesis of Fowler syndrome. **a,** Illustration of treatment regimen for the mice with choline deficient diet (CDD). **b,** Maternal choline deficiency affects embryonic development. Arrow shows failed development of eyes in a WT embryo. **c,** Quantification of body length of WT, HET, and KO E14.5 embryos from normal chow diet (ND) and CDD diet. Each symbol represents one mouse. **d,** Depletion of choline ameliorated the dilation of the CNS blood vessels in the cortical regions but not in the ganglionic eminences (GE) of E14.5 KO embryos. **e,** Diameters of CNS blood vessels in the neocortex of KO embryos were decreased after CDD treatment. Note that there was no improvement in the phenotypes of the blood vessels in the ganglionic eminence (GE) regions. Data are expressed as mean ± SEM. Each symbol represents the averaged diameters of blood vessels from one image. n=3 embryos per genotype. ****P< 0.0001, One-way ANOVA. **f,** Western blot analysis of protein expression for indicated missense mutations. **g,** Immunofluorescent analysis of the localization of the indicated mutant proteins in HEK293 cells. **h,** Missense mutations of Mfsd7c affects choline import activity. Percentage of choline transport activity of the indicated mutants compared to wild-type (WT) Mfsd7c. Data are expressed as mean ± SD. Each symbol represents one replicate.****P< 0.0001, **P< 0.01; One-way ANOVA. Mean percentage of transport activity for each mutant was shown on top of each bar. **j,** Pedigree of the patient’s families. **i,** Sanger sequencing results confirm the mutation P276S from the mother and L483R from the father in a non-consanguinity family. **k,** MRI images of the patient taken at 4 months of age. Arrowhead shows signs of calcification. Arrows show delayed development of cerebellum and neocortex.

Missense mutations of *MFSD7c* have been reported in patients with Fowler syndrome. However, it remains unknown whether these mutations cause a loss or gain of choline function of *MFSD7c*. Thus, we performed mutagenesis to generate several homozygous and compound heterozygous missense mutations including S203Y, T430R, and T430M and tested their transport activity. These homozygous missense mutations S203Y, T430R, and T430M were chosen as the first two former mutations caused lethal Fowler syndrome, whereas the latter mutation caused non-lethal Fowler syndrome ^1, 28, 2,29^. We also examined the activity of missense mutations P340L and T430A that were reported in a non-lethal Fowler patient^1^. First, we confirmed that these missense mutations did not affect the expression and localization of MFSD7c (**Figure 5f-g**). Second, we measured choline transport activity and found that most of these missense mutations of *MFSD7c* exhibited significantly reduced or abolished choline transport activity (**Figure 5h**). A patient homozygous for S203Y was aborted at 20 weeks during gestation^1^. Choline transport activity of this missense mutation was abolished (**Figure 5h**). The T430M mutation was associated with non-lethal Fowler syndrome, whereas homozygous T430R mutation was reported to be lethal^4, 2^. We showed that T430M had a reduction of approximately 62% activity, whereas T430R is an inactive mutant (**Figure 5h**). Previously, compound heterozygous mutants (P340L and T430A) were found in a 2 years-old patient, who is still living^1^. His brain MRI showed mild microcephaly. We found that both alleles had partially decreased choline transport activity (**Figure 5h**). The current study also identified a patient with phenotypes associated with Fowler syndrome. Whole exome sequencing identified two missense mutations of *MFSD7c* coding sequence that lead to a substitution of proline at position 276 to serine (P276S) inherited from the mother and leucine at position 483 to arginine (L483R) from the father (**Figure 5i-k**). The patient exhibits severe microcephaly with neurological disorders (**Figure 5k**). Transport activity of these missense mutants showed that P276S retained approximately 50% choline transport activity, whereas L483R is likely unstable, leading to a severe reduction of activity in MFSD7c (**Figure 5f, h**). Expending these results, we also tested the missense mutations that have been reported in the literature. We found that most of lethal mutations caused a significant reduction of choline transport activity (**Extended Data Figure 9; Supplemental data table 18**). Nevertheless, there were several missense mutations which led to partial loss of choline transport activity, but the patients carrying these mutations exhibited severe clinical features. It is unclear whether other factors also contributed to the pathological conditions in these cases. Together, these results suggest that reduced choline transport activity of MFSD7c may be associated with the pathogenesis of Fowler syndrome. The discovery of the molecular function of MFSD7c as a choline transporter is an important step to facilitate with the understanding of the disease mechanisms in the patients with Fowler syndrome.

## Discussion

The import of choline to the brain via the blood-brain barrier was evidenced several decades ago^18,19,20,30^. Most of these prior studies used *in situ* rat models and reported that exogenous choline is rapidly taken up to the brain. These experiments were often stopped within 1-3 minutes ^18,19,30^. Although these studies hinted that plasma choline is imported to the brain via the BBB, it was rather elusive if choline is taken up to the brain at physiological conditions. To our knowledge, the molecular machinery for choline import at the BBB as anticipated by these prior studies remains uncharacterized. MFSD7c is specifically expressed in endothelial cells of the CNS blood vessels^20,31,32,33^. Deletion of *Mfsd7c* in endothelial cells recapitulates the phenotypes obtained from the global knockout, implicating its major role at the BBB^7^. The current work identifies MFSD7c as a choline transporter, which facilitates choline transport bidirectionally. Employing *Mfsd7c* knockout models, we observe that injection of a high dose of choline to the blood increases choline import to the brain via MFSD7c. This result is consistent with the previous reports and suggests that if a high level of choline is delivered to the blood, a small amount of choline might be imported to the brain by MFSD7c. Nevertheless, the import function of choline by MFSD7c might not be relevant at physiological conditions. Our notion is well in line with numerous reports using radioactive choline (11C or 18F-choline) for cancer diagnosis. Most of radioactive choline is accumulated in the peripheral organs such as liver, kidney, and prostate ^34,35,36^. Unexpectedly, our results show that MFSD7c at the CNS blood vessels is required for the export of choline from the brain parenchyma. We demonstrate that lysophosphatidylcholine (LPC) delivers choline in the brain. The remodelling of phospholipids generated from plasma derived LPC liberates choline in the brain which is expected to be imported back to the CNS endothelial cells by MFSD7c. We argue that LPC provides a sufficient amount of choline in the brain. The levels of plasma LPC can reach to 400-500µM ^25^, whereas plasma choline level is approximately 10-15µM ^22,23^. Plasma LPC is transported to the brain by Mfsd2a which is one of the highest genes expressed in the BBB^37,38,26,32^. Thus, we argue that LPC provides a major source of choline to the brain, not free choline from the plasma. Remarkably, our stable isotopic tracing results reveal that there is a rapid remodelling of LPC/PC in the brain, which liberate free choline. The excessive amount of choline is likely to be taken back to the endothelial cells by Mfsd7c. Consistently, deletion of *Mfsd7c* results in increased levels of endogenous choline in the brain. Thus, it is likely that the major physiological function of MFSD7c at the BBB is to reduce parenchymal choline levels in the brain. At this point, it is still unclear if MFSD7c exports excessive brain choline to the plasma or choline is accumulated in the endothelial cells. This remains to be investigated. Nevertheless, the increased levels of acetylcholine in the brain of Mfsd7c knockout mice might hint that choline is indeed accumulated in the brain parenchyma since acetylcholine is mainly generated by cholinergic neurons ^39^. Furthermore, we show that missense mutations of *MFSD7c* result in reduced choline transport activity. Based on our results, we propose that accumulation of choline in the brain due to Mfsd7c mutations might be a casual factor for Fowler patients. Our work might provide a potential mechanism linking the defective choline export functions of MFSD7c to the pathogenesis of Fowler syndrome. In summary, the discovery of MFSD7c as a choline transporter at the BBB provides a foundation for future research to reveal the disease mechanism of Fowler syndrome as well the metabolic fates of choline in the brain.

## Material and methods

### Mice

Global deletion of *Mfsd7c* has been described previously ^1^. To generate the postnatal deletion of *Mfsd7c* in the blood brain barrier, *Mfsd7c*^f/f^ was crossed with Cdh5-CreERT^2^ mice ^40^ to generate *Mfsd7c*^f/f^, Cdh5-CreERT2 mice (Ec*Mfsd7c*-KO). Male and female mice with age >7-8 weeks old were used for experiments. Mice were maintained at a constant temperature of 20°C with 12-hour light/12-hour dark cycle on normal chow diets. All experimental protocols and procedures in the protocol R19-0567 were approved by IACUC committees under National University of Singapore.

### Plasmids

All plasmids were constructed in pcDNA3.1 vector for overexpression unless stated otherwise. hChKA (D10704.1), ETNK1, CHAT (NM_020986.4) gene were synthesized from GenScript and inserted into pcDNA3.1. pcDNA3.1hMfsd7c, pcDNA3.1hMfsd7cT430M and S203Y were generated previously^1^. The other missense mutations including hMfsd7cT430A, P276S, L483R, P340L, and T430R were generated by mutagenesis and confirmed by Sanger sequencing. Zebrafish (DaMfsd7c_a, XM_688497.7; DaMfsd7c_b: XM_003200755.5), medaka fish (MeMfsd7c, XM_004082328.4), and frog Mfsd7c (XeMfsd7c, NM_001016982.2) coding sequences were synthesized by GenScript and cloned into pcDNA3.1 plasmid for overexpression. The pESC-HIS-hMfsd7c plasmid (Genscript) was used for overexpression on the yeast *S. cerevisiae*.

### Untargeted metabolite analysis

Untargeted metabolomics analysis was performed at the Genome Analysis Center, Research Unit Molecular Endocrinology and Metabolism, Helmholtz Center Munich. Frozen WT and *Mfsd7c* knockout embryonic brain and liver samples at E14.5 were weighed and placed into pre-cooled (dry ice) 2 ml homogenization tubes containing ceramic beads with a diameter of 1.4 mm (Precellys® Keramik-Kit 1.4 mm). Pre-cooled water with a ratio of 5 µl/mg tissue was added into the tubes. The samples were then homogenized in Precellys 24 homogenizer (PEQLAB Biotechnology GmbH, Germany) equipped with an integrated cooling unit for 3 times 20 s at 5,500 rpm, with 30 second intervals (to ensure freezing temperatures in sample vials) between the homogenization steps. A hundred µl of the brain and liver homogenates were loaded onto two separate 2 ml 96-deep well plate, one plate for the brain sample set and the other plate for the liver samples. Three types of quality control samples were analyzed in concert with the experimental samples in each sample set: samples generated from a pool of human plasma, samples generated from a small portion of each experimental sample from the respective sample set served as technical replicate throughout the data set; and extracted water samples served as process blanks. Experimental samples and controls were randomized across the metabolomics analysis.

The homogenates in each well of the 2 ml 96-deep well plate were extracted with 500 µl methanol, containing four recovery standards to monitor the extraction efficiency. After centrifugation, the supernatant was split into aliquots in two 96-well microplates. The first 2 aliquots of each sample set were used for ultra-high performance liquid chromatography-tandem mass spectrometry (UPLC-MS/MS) analysis in positive electrospray ionization modes (i.e., early and late eluting compounds), 1 aliquot of each sample set was used for UPLC-MS/MS analysis in negative ionization mode, and the rest were kept as reserve for backup. The extract aliquots were dried on a TurboVap 96 (Zymark).

Prior to UPLC-MS/MS analysis, the dried extract samples were reconstituted in acidic or basic LC-compatible solvents, each of which contained 8 or more standard compounds at fixed concentrations to ensure injection and chromatographic consistency. The UPLC-MS/MS platform utilized a Waters Acquity UPLC with Waters UPLC BEH C18-2.1×100 mm, 1.7 μm columns and a Thermo Scientific Q-Exactive high resolution/accurate mass spectrometer interfaced with a heated electrospray ionization (HESI-II) source and Orbitrap mass analyzer operated at 35,000 mass resolution. Extracts reconstituted in acidic conditions were gradient eluted using water and methanol containing 0.1% formic acid, while the basic extracts, which also used water/methanol, contained 6.5mM ammonium bicarbonate. The MS analysis alternated between MS and data-dependent MS2 scans using dynamic exclusion, and the scan range was from 80-1000 m/z.

Metabolites were identified by automated comparison of the ion features in the experimental samples to a reference library of chemical standard entries that included retention time, molecular weight (m/z), preferred adducts, and in-source fragments as well as associated MS spectra and curated by visual inspection for quality control using software developed at Metabolon. Chromatographic peaks were quantified using area-under-the-curve.

### *In vivo* transport of radiolabelled choline

To study the uptake of choline into the brain, control (*Mfsd7c*^f/f^) and EcMfsd7c-KO mice aged 4-6 months was intravenously injected 2mM radiolabelled choline (stock: 333mM choline with 0.05µCi/µl). After 5 mins of injection, 50µl blood was collected and the radioactive levels were quantified to monitor the initial radiolabelled levels in the circulation. After 30 mins of injection, mice were perfused with 20ml of PBS. Then, brains, lungs, hearts, kidneys, and livers were collected for quantification of radioactive signals. Similar amount of tissues was homogenized in RIPA buffer and the radioactive signals were quantified by liquid scintillation counter. The radioactive signals from tissue from each mouse were normalized to radioactive signals from blood collected at 5 mins post-injection.

### Choline-d9 and LPC-d49 experiments

For stable isotopic tracing with choline-d9 (containing 9 deuterium atoms) injection, pregnant Mfsd7c+/− mice at 12.5 days of gestation were injected daily by IV route with an amount of 2mM choline-d9/day until gestation day 14.5 (total of 3 doses of choline-d9 from E12.5-14.5). The embryos were collected at E15.5 after the last dose at E14.5 and genotyped. The brains and livers of embryos were harvested for lipid extraction for lipidomic analysis. Briefly, equal amount of brain and liver lysates from WT, HET and KO embryos were extracted with 1-butanol:methanol (ratio 1:1) method for choline and lipid analyses by LC-MS/MS.

For stable isotopic LPC-d49 (Sigma: 870308P-5MG; containing choline-d13, glycerol-d5, and palmitate-d31) injection, Ec*Mfsd7c*-KO and control mice aged 7-10 months were injected with a dose of 300µM LPC-d49 in 12% BSA. After 2 hours of injection, plasma was collected from blood; then the mice were perfused with cold PBS to remove blood. After that the brains and livers were collected. Brains and livers were homogenized in PBS and the same amount of tissue lysates was used for lipid and choline extraction with chloroform:methanol (1:2 v/v) method ^42^. Organic phase was used for lipid analysis and aqueous phase (upper phase) was used for choline analysis by LC-MS/MS.

### Lipids and choline quantification by LC/MSMS

Samples were randomized before extraction and analysis. Blank samples (10 µL MilliQ water), matrix blanks (samples extracted without spiking internal standards, see below), and pooled QC samples were used to assess method performance. In addition, diluted pooled QC samples were used to assess response linearity. Blanks, blank extracts, QC, and diluted QC were interspersed with study samples throughout the analytical run. Phospholipids and choline were separated using HILIC column (Kinetex 2.6 µm HILIC 100 Å, 150 × 2.1 mm, Phenomenex, Torrance, CA, USA) on Agilent 6495A and 6495C triple quadrupole mass spectrometers (Agilent Technologies). Mobile phases A (50% acetonitrile (LC-MS grade, Thermo Fisher Scientific Inc.) 50% 25 mM ammonium formate (Sigma-Aldrich) adjusted to pH = 3.5) and B (95% acetonitrile and 5% of 25 mM ammonium formate adjusted to pH = 3.5) were mixed at the following gradient: 0–6 min, 99–75% B; 6–7 min, 75–10% B; 7–7.1 min, 10–99% B; 7.1–10.1 min, 99% B. The flow rate was 0.6 mL/min, and the sample injection volume was 1 µL. MS parameters were as follows: electrospray ionization, gas temperature 200°C, gas flow 12 L/minute, sheath gas flow 12 L/minute, and capillary voltage 3,500 V. Phospholipids were quantified at the sum composition level using multiple reaction monitoring (MRM) using transition to phosphocholine headgroup (*m/z* 184 for endogenous lipids and adjusted to appropriate *m/z* for deuterated lipids). Choline was quantified using multiple reaction monitoring (MRM) using transition 104 to 60 (adjusted to appropriate *m/z* for deuterated choline). These internal standards (IS) were used for lipidomics and choline analysis by LC MS/MS. Internal standards solutions were prepared in butanol/methanol (BuMe) (1:1, v/v) or chloroform/methanol (1:2) containing phospholipid IS SPLASH Mix (Avanti, 330707) (containing 5.3 µM 15:0-18:1(d7) PC, 0.19 µM 15:0-18:1(d7) PE, 0.13 µM 15:0-18:1(d7) PS (Na Salt), 0.93 µM 15:0-18:1(d7) PG (Na Salt), 0.27µM 15:0-18:1(d7) PI (NH4 Salt), 0.27 µM 15:0-18:1(d7) PA (Na Salt), 1.20 µM 18:1(d7) Lyso PC, 0.27 µM18:1(d7) Lyso PE, 13.32 µM18:1(d7) d18:1-18:1(d9) SM. Choline-d9 was used as IS in choline analysis.

### Transport assays in HEK293 cells

Human (hMfsd7c) or mouse (mMfsd7c) Mfsd7c cDNA constructed in pcDNA3.1 was transfected to HEK293 cells using lipofectamine 2000. Mock was treated with empty plasmid. For experiments in which human choline kinase A (hChKA) was used, 1-1.5µg pcDNA3.1 hChKA plasmid was co-transfected with 2-2.5µg plasmid of hMfsd7c or mutant plasmids. Cells transfected with 1-1.5µg pcDNA3.1 hChKA plasmid was used as control. After 18-30 hours post-transfection, cells were used for transport assays with 100µM [^3^H] choline (ARC: ART 0197) in DMEM containing 10%FBS. The transport assays were stopped after 1hour incubation at 37°C by washing once with cold plain DMEM medium. Cell pellets were lyzed in RIPA buffer and transferred to scintillation vials for quantification of radioactive signal using Tricarb liquid scintillation counter. Transport activity of MFSD7c was expressed as DMP for [^3^H] isotopes and CPM [^14^C] isotopes. For dose dependent transport activity, HEK293 cells were similarly transfected with hMfsd7c or hMfsd7cS203Y mutant plasmid alone or co-transfected with hChKA. Transport activity was conducted with 20, 45, 90, and 250 µM [^3^H] choline in DMEM containing 10% FBS for 1 hr at 37^0^C. For time-dependent assay, hMfsd7c was transfected or co-transfected with hChKA. Transfected HEK293 cells were incubated with 100µM [^3^H] choline in DMEM containing 10%FBS for 15, 30, 60, and 120 mins at 37^0^C. For transport assay with [^14^C] ethanolamine, [^3^H] L-carnitine, and [^3^H] acetyl-carnitine, [^3^H] palmitoyl-carnitine, [^14^C] octanoyl-carnitine, HEK293 cells were co-transfected with *MFSD7c* and hChKA as described above. Transport assays were performed with 100µM [^14^C] ethanolamine, 1mM [^3^H] L-carnitine, 1mM [^3^H] acetyl-carnitine, 10µM [^3^H] palmitoyl-carnitine, or 100µM [^14^C] octanoyl-carnitine in DMEM with 10%FBS for 1 hour. Cells were washed once with cold plain DMEM and lyzed with RIPA buffer for radioactive quantification. For import assays with the missense mutants, these plasmids were co-transfected with hChKA to HEK293 cells. Import assays were performed with 100µM [^3^H] choline in DMEM containing 10% FBS for 60 mins. Radioactive signals from cells were quantified and converted to percentage of WT activity.

For export assays, HEK293 cells were transfected with empty, hMfsd7c, or mMfsd7c plasmids. After 24 hours of transfection, cells were loaded with 500µM [^3^H] choline for 1 hour. The cells were then washed to remove remaining [^3^H] choline in the medium and incubated with DMEM medium to stimulate the release. The cells were harvested at 0 (before release) and 1hour (after release) after incubation with DMEM for radioactive quantification as described above.

### Sodium and pH-dependent transport assays

For sodium dependent assays, sodium is replaced with lithium in transport buffer (NaCl buffer: 150mM NaCl, 5mM KCl, 10mM HEPES-Na, pH 7.4; LiCl buffer: 150mM LiCl, 5mM KCl, 10mM HEPES, pH 7.4 with HCl). For pH dependent assays, NaCl buffer was adjusted to pH 6.5 and pH 8.5. For transport assay conditions, overnight *hMfsd7c* and *hChKA* co-transfected HEK293 cells were washed once with the same buffer before incubated with 100µM [^3^H] choline for 15 minutes at 37°C. Cells were lyzed in RIPA buffer and mixed with scintillation fluid for radioactive quantification.

### Electrophysiology

For single cell recording of membrane potential alteration during transport of choline, HEK293FT cells were cultured in DMEM medium supplemented with 10% fetal bovine serum and 1% penicillin and streptomycin and maintained in 5% CO_2_ incubator at 37°C. For transfection, cells were seeded on the Petri dishes and grown overnight. Subsequently, 2 µg of empty pIRES2-EGFP (empty plasmid), pIRES2-EGFP *hMfsd7c* or mutant pIRES2-EGFP S203Y plasmid was co-transfected with 2 µg of pIRES2-RFP *hChKA* plasmid using lipofectamine 2000. The transfected cells were incubated for 24 h in 5% CO_2_ incubator at 37°C. At 48 hours post-transfection, the cells were split and seeded on poly-D lysine coated cover slips for one more day before recording.

### Whole-cell patch-clamp recordings and data analysis

To record the changes in resting membrane potential, the internal solution (pipette solution) contained 130mM K-gluconate, 10nM KCl, 5mM EGTA, 10mM HEPES, 1mM MgCl_2_, 0.5mM Na_3_GTP, 4mM Mg-ATP, 10mM Na-phoshocreatine, pH 7.4 (adjusted with KOH) was filled in the pipette tip. The external solution contained: 10mM Glucose, 125mM NaCl, 25mM NaHCO_3_, 1.25mM NaH_2_PO4.2H20, 2.5mM KCl, 1.8mM CaCl_2_, 1mM MgCl_2_, pH 7.4 (300–310 mOsm). Single cells with both EGFP (empty, *hMfsd7c* or *S203Y* plasmid) and RFP (*hChKA*) fluorescence was visualized under fluorescent microscope for patching. Resting membrane potential was recorded under current clamp with an Axopatch200B or multiclamp 200B amplifier (Molecular Device) with 0 pA current injection. Subsequently, 200 µM choline was perfused to the cells after 2 minutes of stable base line recording. After 5 min of recording, the ligand was washed away. Approximately, 10 individual cells were recorded for each condition. The data were analyzed using Graphpad Prism V software (San Diego, CA) and Microsoft (Seattle, WA) Excel. Data are expressed as mean values ± SEM.

### Expression of human *Mfsd7c* in *S. cerevisiae* Hnm1 mutants

*Saccharomyces cerevisiae* acquires choline via HNM1. The *S. cerevisiae Hnm1* mutants were acquired from Dharmacons. Wildtype and *Hnm1* mutant yeasts were grown in Yeast extract-Peptone-Dextrose (YPD) medium at 30^0^C overnight before transformation. The pESC-HIS-*hMfsd7c* plasmid was transformed into wildtype and *Hnm1* mutant yeasts following the previously described protocol ^43^. After heat shock at 42^0^C for 30 minutes and recovery in YPD for 2 hours, the yeasts were spread on 2% agar plates prepared in yeast nitrogen base (YNB) medium (Sigma Aldrich) supplemented with 2% glucose and yeast synthetic drop-out medium supplements without histidine (Sigma Aldrich). Positive colonies were picked up randomly and further confirmed by transport assay and western blot.

### Transport assay for yeast cells

Wild type, *Hnm1* mutant and transformed yeast cells were grown in YNB medium with 2% glucose and drop-out medium supplements without histidine at 30^0^C overnight followed by induction of hMfsd7c expression in YNB medium with 2% galactose for at least 5 hours at 30^0^C. An 800µl of yeast cells at OD_600_=1 was centrifuged, washed twice with distilled water, then incubated in PBS containing 100 µM [^3^H] choline for 10 mins. The transport assay was stopped by washing twice with PBS containing 100 µM choline. Yeast cells were lyzed with lysis buffer containing 2% Triton-X, 1% SDS, 100mM NaCl, 10mM Tris-HCl (pH 8.0), 1mM EDTA (pH8.0) at 95^0^C for 5 mins for radioactive quantification by liquid scintillation counter.

### Western blot analysis

For protein extraction, transfected HEK293 cells with *hMfsd7c*, *mMfsd7c*, *hMfsd7c* mutants were lysed with RIPA buffer. For validation of deletion of *Mfsd7c* in mice, micro-vessels from PBS perfused brains of WT and Ec*Mfsd7c*-KO mice were isolated, homogenized and lyzed with RIPA buffer. For Western blot analysis of hMfsd7c expression in yeasts, 1ml of yeast culture at OD_600_=1 from WT, *Hnm1* mutant yeasts or transformed yeasts was used for total protein extraction. BCA assay was used for total protein quantification. These antibodies were used: VDAC (CellSignaling CS4866), CoxIV (Invitrogen, MA5-15686), OPA1 (BD biosciences, 612602), MRPS35 (Protein biotech, 16457), CHKA (CellSignaling, CS13422S), HMOX1 (Proteintech, 10701-1-AP), NDUFS1 (Proteintech, 12444-1-AP). Western blot analysis was conducted as previously described^1^.

### Fowler patient recruitment

A five-month-old infant born at 39w1d with multifocal epilepsy, ventriculomegaly, developmental delay, and lissencephaly was identified via GeneMatcher collaboration ^44^. She was born to a 38 years old mother. Pregnancy was complicated by gestational diabetes and maternal hypertension. There was concern for lissencephaly on prenatal ultrasound and MRI. Prenatal CMA and karyotype were normal. She was delivered vaginally complicated by shoulder dystocia. She required brief positive pressure ventilation and was admitted to the NICU on NIPPV. Apgars at 1 and 5 minutes were 4 and 9 respectively. Birth parameters include length of 49 cm (47th percentile WHO), weight of 2.86 kg (20th percentile WHO) and head circumference of 32 cm (6th percentile WHO). Her whole exome sequencing at birth showed two variants of uncertain significance in FLVCR2, c.826C>T (Maternal) and c.1448T>G (Paternal). The missense mutations were confirmed by Sanger sequencing. There was no family history of a similar presentation. This was the first pregnancy between this patient’s parents. Mother had a previous miscarriage and a healthy older daughter. Parents report no consanguinity.

### RNA sequencing and analysis

For bulk RNA sequencing of whole brains from E14.5 wildtype and KO embryos from normal chow diet (ND), this has been deposited in GEO database with the accession number: GSE148854. For RNA sequencing of isolated endothelial cells from heterozygous (HET) and KO embryos at E14.5, we re-analyzed the dataset GSE129838 ^7^. For RNA sequencing of whole brains from E14.5 wildtype embryos from choline deficient diet (CDD), pregnant wildtype mice were fed with CDD diet at the time-mating day until embryo collection. The whole brain of E14.5 embryos was collected for bulk RNA sequencing. This dataset has been deposited in the GEO database with the following accession number: GSE239589.

### Immunofluorescence staining for neocortical vessels

*Mfsd7c^+/−^* female mice were fed on choline-deficient diet for 2 weeks and then time-mated with the heterozygous male. E14.5 embryos were dissected from the pregnant female for collection of brains for immunostaining of CNS vessel morphology. Brain section was permeabilized in 0.5% triton-100 (PBST), blocked in 5% normal goat serum and then incubated with anti-mouse GLUT1 (Abcam, ab40084,1:200) at 4°C overnight, followed by 3 washes in PBS at 10-minute intervals. Then, the goat anti-mouse antibody (Alexa Fluor 488, Thermofisher A11034,1:500) was added to the slides for 1-hour incubation at room temperature to visualize the GLUT1 signal. Images were taken by the confocal microscope Zeiss LSM710. Vessel phenotypes were analysed using measurement tools in imageJ.

### Choline and acetylcholine assays

Choline and acetylcholine from plasma and brain samples were measured by fluorometric assay (Abcam: ab65345). Tissues were homogenized in choline buffer. Choline and acetylcholine were assayed according to the manufacturer’s instructions.

### Transmission electron microscope

Wild-type (n=3) and *Mfsd7c^−/−^* (n=3) embryos at E14.5 were collected from the same pregnant mice and fixed in 2.5% glutaraldehyde for 24 hours at room temperature. Brain sections were prepared and were post-fixed with 1% osmium tetroxide for 1 hour and dehydrated using a serial ethanol concentration. Images were taken using a Fei Quanta650 transmission electron microscope.

### Statistical analysis

Data was analyzed using GraphPrism9 software. Statistical significance was calculated using parametric and non-parametric *t*-tests, One or Two-way ANOVA as indicated in the figure legends. P value < 0.05 was considered as statistically significant.

## Data Availability

All data have been included in the manuscript.

## Author Contributions

L.N.N., T.N.U.L., and T.Q.N. and performed *in vitro* transport assays; T.Q.N. and L.N.N. performed *in vivo* transport assays. A.A. and J.A. performed metabolite analysis. D.Y., H.H., and T.W.S. performed electrophysiological assays. Q.H., F.L, helped with cell cultures. T.N.U.L., A.V.S., and M.C. performed yeast experiments. H.T.T.H., X.T.A.N., T.Q.N., A. C-G., P.Y.L., and M.W. performed lipidomics. A.L. and C.G. provided clinical data. D.T.N and N.C.P.L. performed transport assays and Western blot for mutants. X.T.A.N. performed in vivo experiments. L.N.N. conceived and designed the study and experiments; analyzed data, prepared the figures, and wrote the paper.

## Acknowledgements

This study was supported in part by Singapore Ministry of Health’s National Research Council NMRC/OFIRG/0066/20, Ministry of Education MOE2018-T2-1-126, NUSMED-FOS Joint Research Programme on Healthy Brain Aging grants (to L.N.N.). The Synthetic Biology Translational Research Programme of the Yong Loo Lin School of Medicine of the National University of Singapore (NUHSRO/2020/077/MSC/02/SB) (to M.W.C.). We thank L.K.V. for providing technical assistance with SLC44a1/2.

---

**Extended Data Fig. 1.**
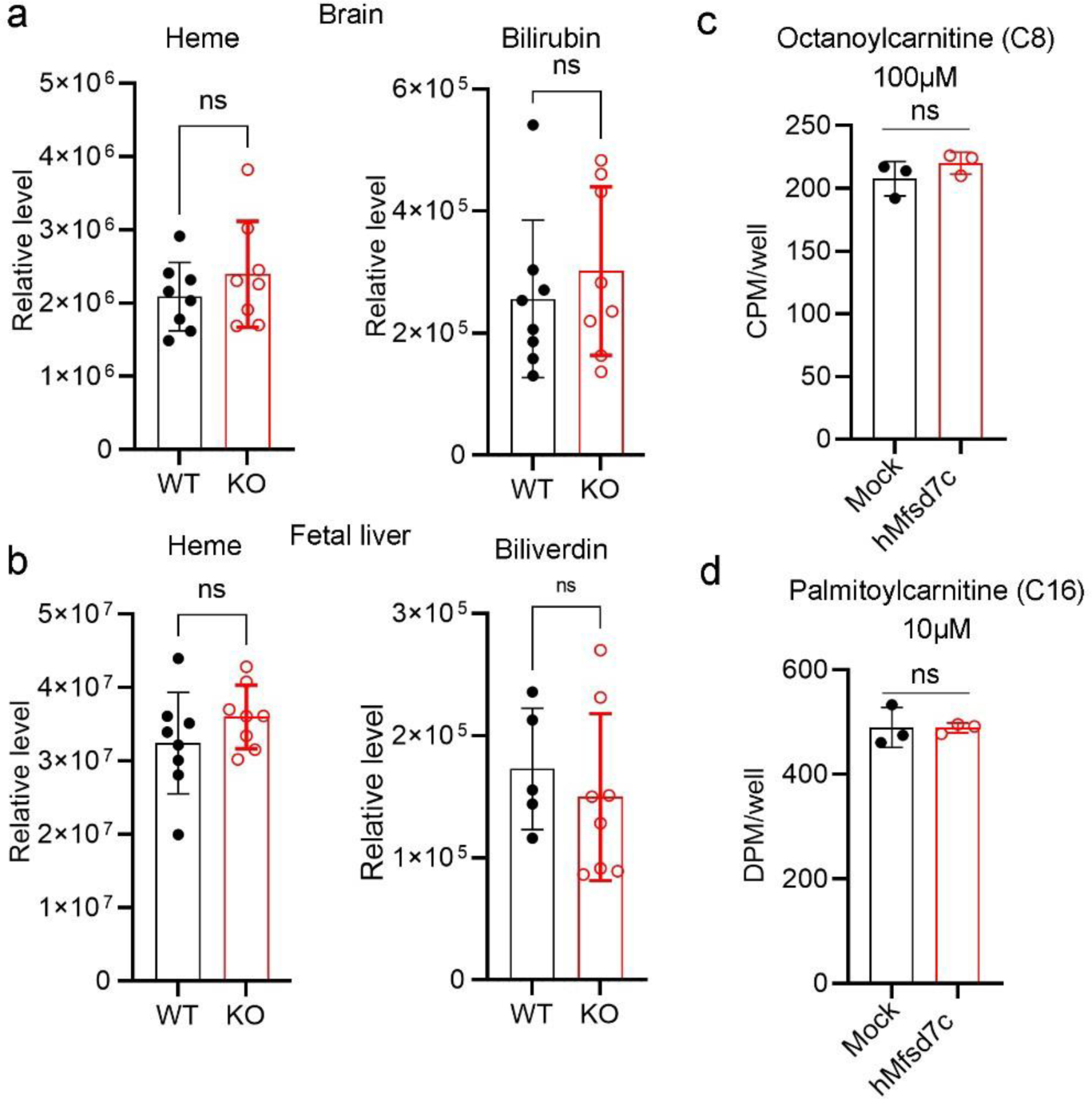
Comprehensive metabolite analysis did not detect changes in heme and heme metabolites in the brain of Mfsd7c knockout, related to Figure 1. **a,** heme and bilirubin levels in the brains of WT and Mfsd7c^−/−^ (KO) embryos. **b,** heme and biliverdin levels in the livers of WT and Mfsd7c^−/−^ (KO) embryos. Each symbol represents on embryo. ns, not significant. **c-d,** Import assays for octanoylcarnitine and palmitoylcarnitine in HEK239 cells overexpressed with hMfsd7c. Mfsd7c did not import these long chain acyl-carnitines. Experiments were repeated thrice in triplicate. ns, not significant. t-test was used. Full list of metabolites can be found in the **Supplemental data tables 1-10**.

**Extended Data Fig. 2.**
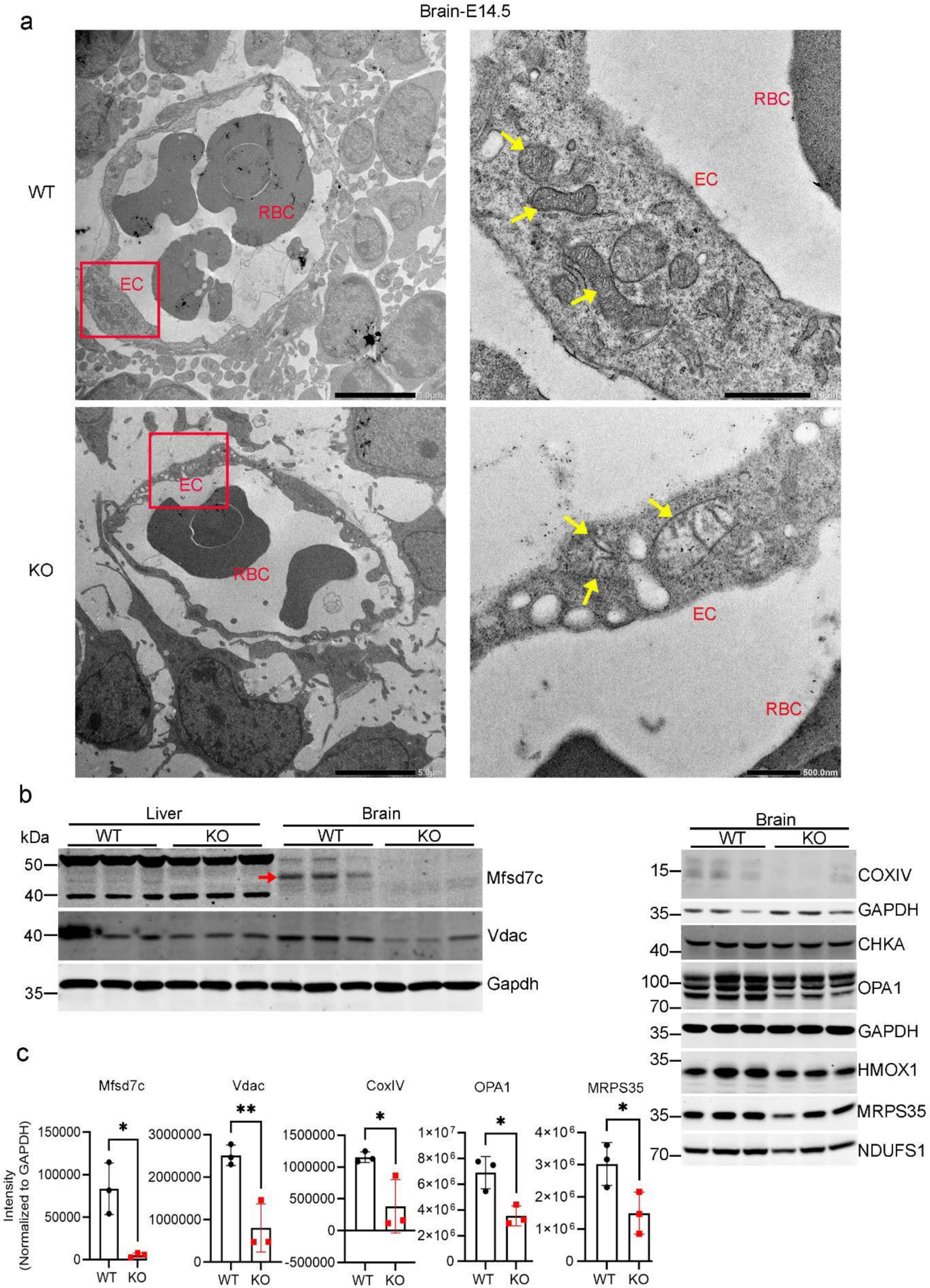
Mitochondrial defects in the brain of Mfsd7c knockout embryos. **a,** Representative images of electron microscopic graphs of mitochondrial morphology of the CNS endothelial cells from E14.5 Mfsd7c KO and controls. Arrows show mitochondria which were enlarged in the Mfsd7c knockouts. n=3 per genotype. **b**, Western blot analysis of Mfsd7c expression in the livers and brains as well as the mitochondrial markers (CoxIV, OPA1, MRPS35, VDAC, NDUFS1) cytosolic markers such as choline kinase A (CHKA) and heme oxygenase 1 (Hmox1) in the brains of E14.5 Mfsd7c KO and controls. Mfsd7c expression is present in the brain, not liver of the embryos. **c**, Quantification of the mitochondrial protein bands from the Western blot analysis shown in b. The levels of mitochondrial proteins were significantly reduced in the brains of E14.5 Mfsd7c KO compared to controls. Experiments were repeated twice in triplicate (n= 3 per genotype). **P<0.01, *P<0.05; ns, not significant. t-test was used.

**Extended Data Fig. 3.**
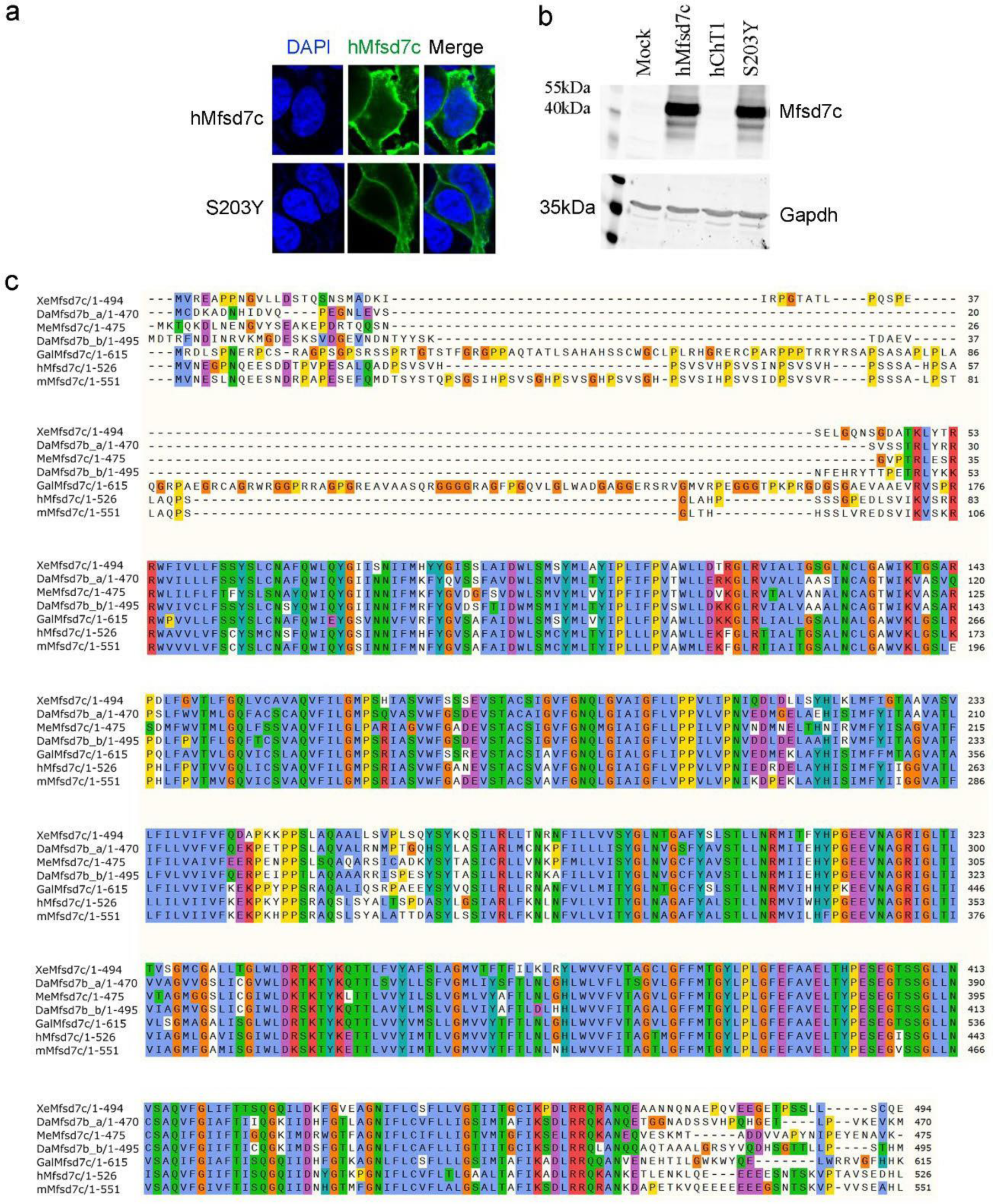
Mfsd7c is localized in the plasma membrane and conserved from fish to man, related to Figure 2. **a-b,** Expression of hMfsd7c and S203Y mutant in HEK293 cells. hChT1 is neuronal choline transporter was used as negative control for Western blot analysis. **c**, Alignment of human (hMfsd7c), frog (XeMfsd7c), Zebrafish (DaMfsd7c_a and DaMfsd7c_b isoforms), Medaka fish (mMfsd7c), Chicken (GaMfsd7c), and mouse (mMfsd7c) Mfsd7c orthologs. Mfsd7c is not conserved in yeasts.

**Extended Data Fig. 4.**
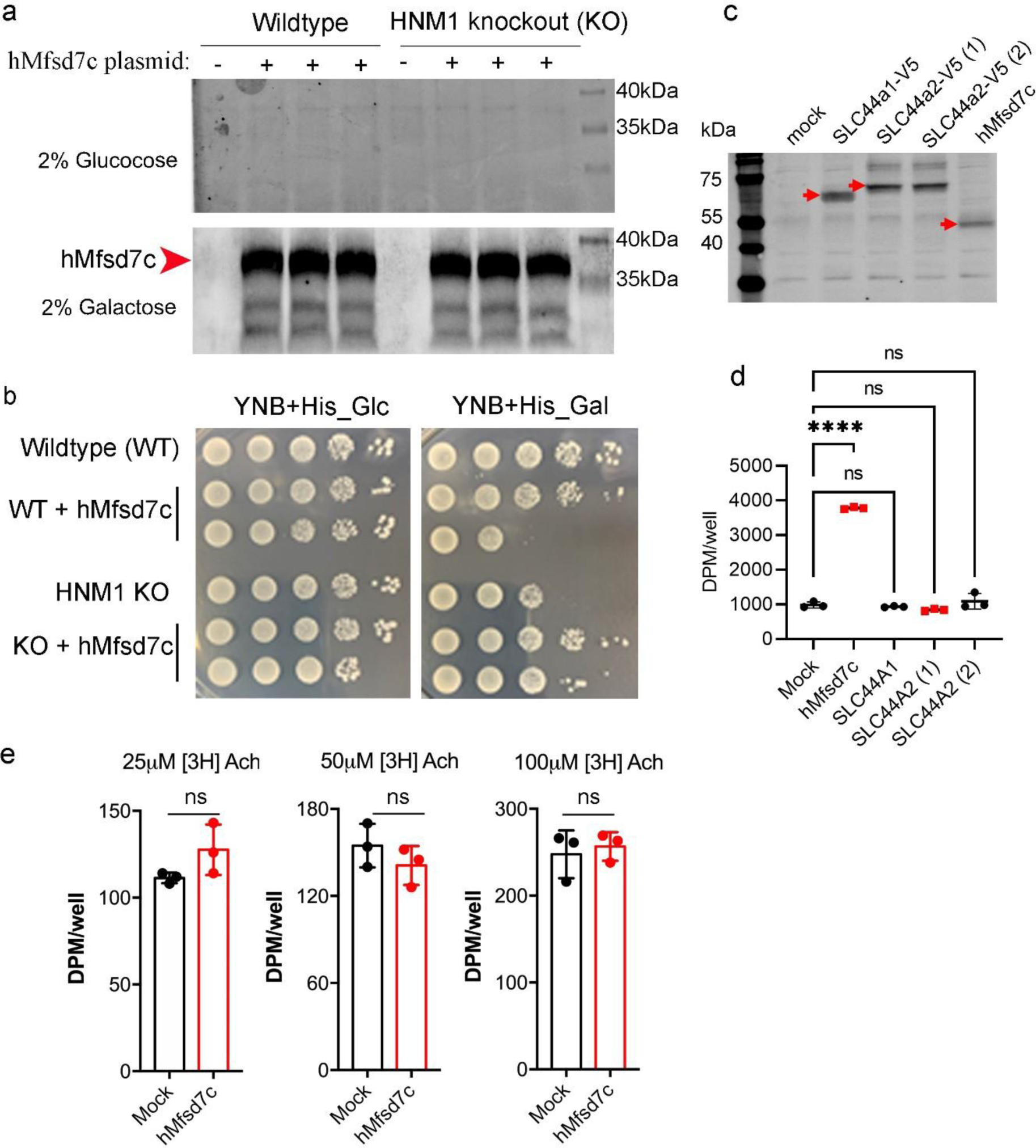
Human Mfsd7c functionally rescues for the loss of HNM1, sole the choline transporter in yeasts. **a-b**, Overexpression of human Mfsd7c in wild-type and HNM1 knockout yeast cells. Gal1 promoter was used for expression of hMfsd7c. Thus, hMfsd7c expression is repressed by glucose (Glc) and induced by galactose (Gal). Yeasts with overexpression of hMfsd7c grew normally in YNB medium with glucose or galactose. Note that 2-3 different transformants with overexpression of hMfsd7c were used. Experiments were repeated with reproducibility. **c**-**d.** human SLC44a1 and SLC44a2 did not exhibit choline import activity. Human SLC44a1-V5 and SLC44a2-V5 were used. **e**, Mfsd7c does not transport acetylcholine. ****P<0.0001, One-way ANOVA was used in d; t-test was used in e.

**Extended Data Fig. 5.**
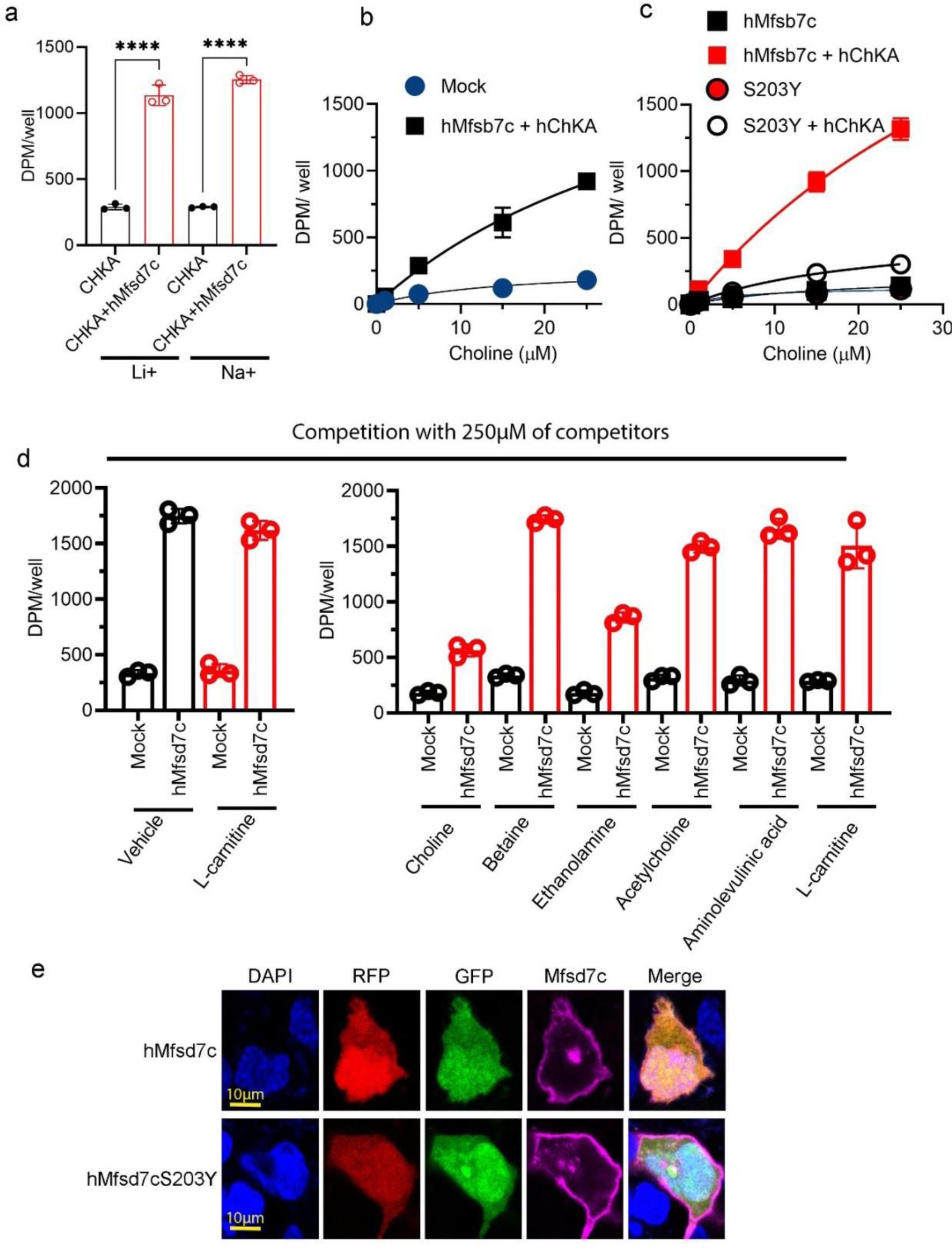
Choline transport properties of Mfsd7c, related to Figure 3. **a**, Mfsd7c activity is not sodium dependent. In these experiments, Mfsd7c was co-expressed with choline kinase A (hChKA). Transport assays were performed in transport buffer with lithium (Li+) in replacement of sodium (Na+). ****P<0.0001. One-way ANOVA was used. **b-c**, Dose-curve of choline transport activity by hMfsd7c and S203Y mutant. Experiments were stopped after 30 mins of choline incubation. Experiments were performed in triplicate. **d**, Competition assays of choline with indicated compounds. A 5-fold excess (250µM) of cold choline and ethanolamine, but not other indicated compounds reduced the import of radioactive choline [^3^H]-choline. Experiments were repeated twice in triplicate. Data are expressed as mean ± SD. **e**, Representative images of co-expression of CHKA in RFP plasmid with hMfsd7c in GFP plasmid for single cell patch clamp. Cells with co-expression of the two fluorescent proteins were selected for patch clamp. Note that Mfsd7c (magenta) is localized in the plasma membrane.

**Extended Data Fig. 6.**
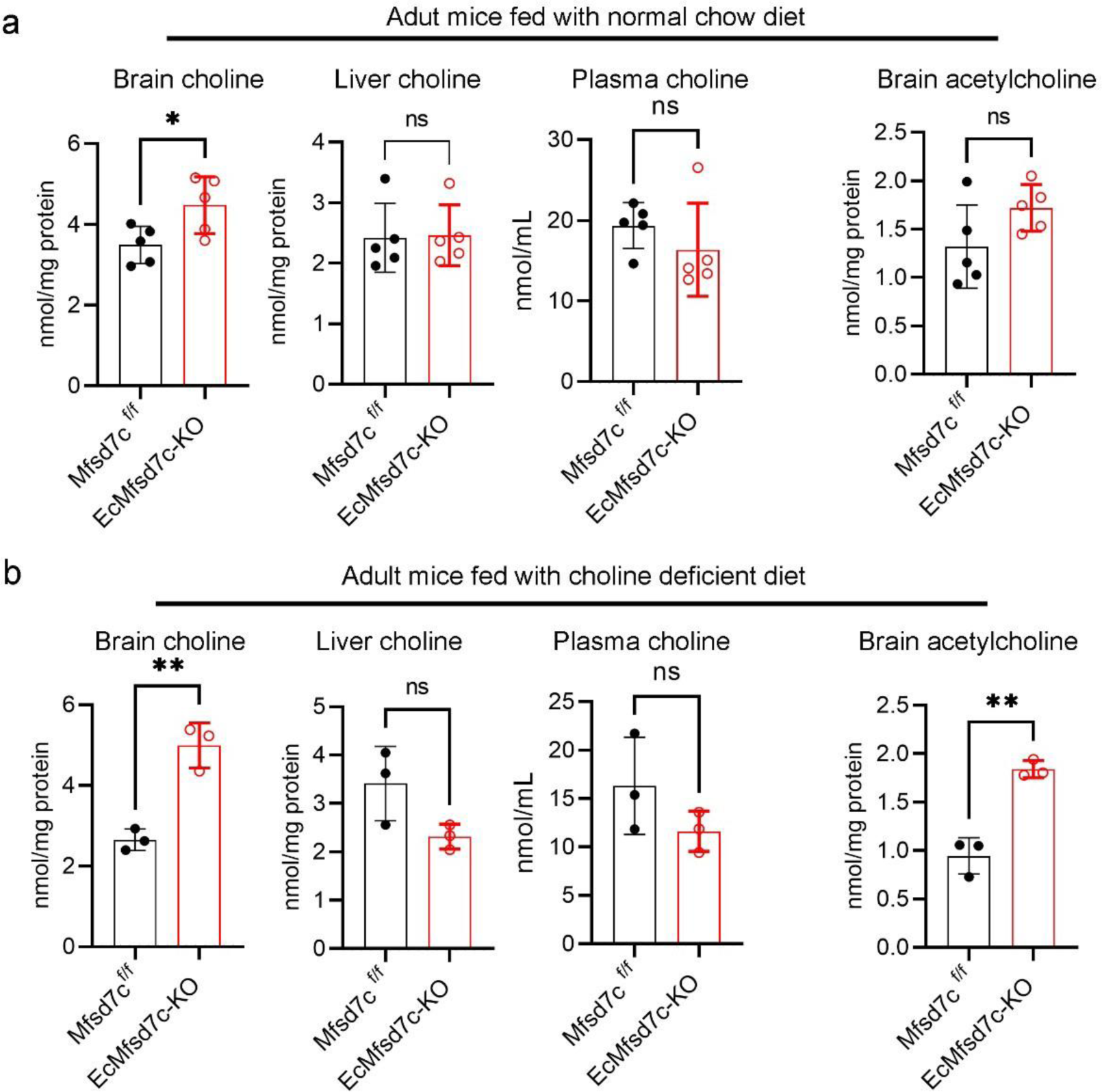
Increased levels of choline and acetylcholine in the brain of adult EcMfsd7c-KO mice. **a,** choline and acetylcholine levels in the brain, liver, and plasma of EcMfsd7c-KO and control mice under normal chow diet. **b,** choline and acetylcholine levels in the brain, liver, and plasma of EcMfsd7c-KO and control mice after feeding for 1-1.5months with choline deficient diet. Each symbol represents one mouse. Data are expressed as mean ± SD. **P<0.001, *P<0.05; t-test was used.

**Extended Data Fig. 7.**
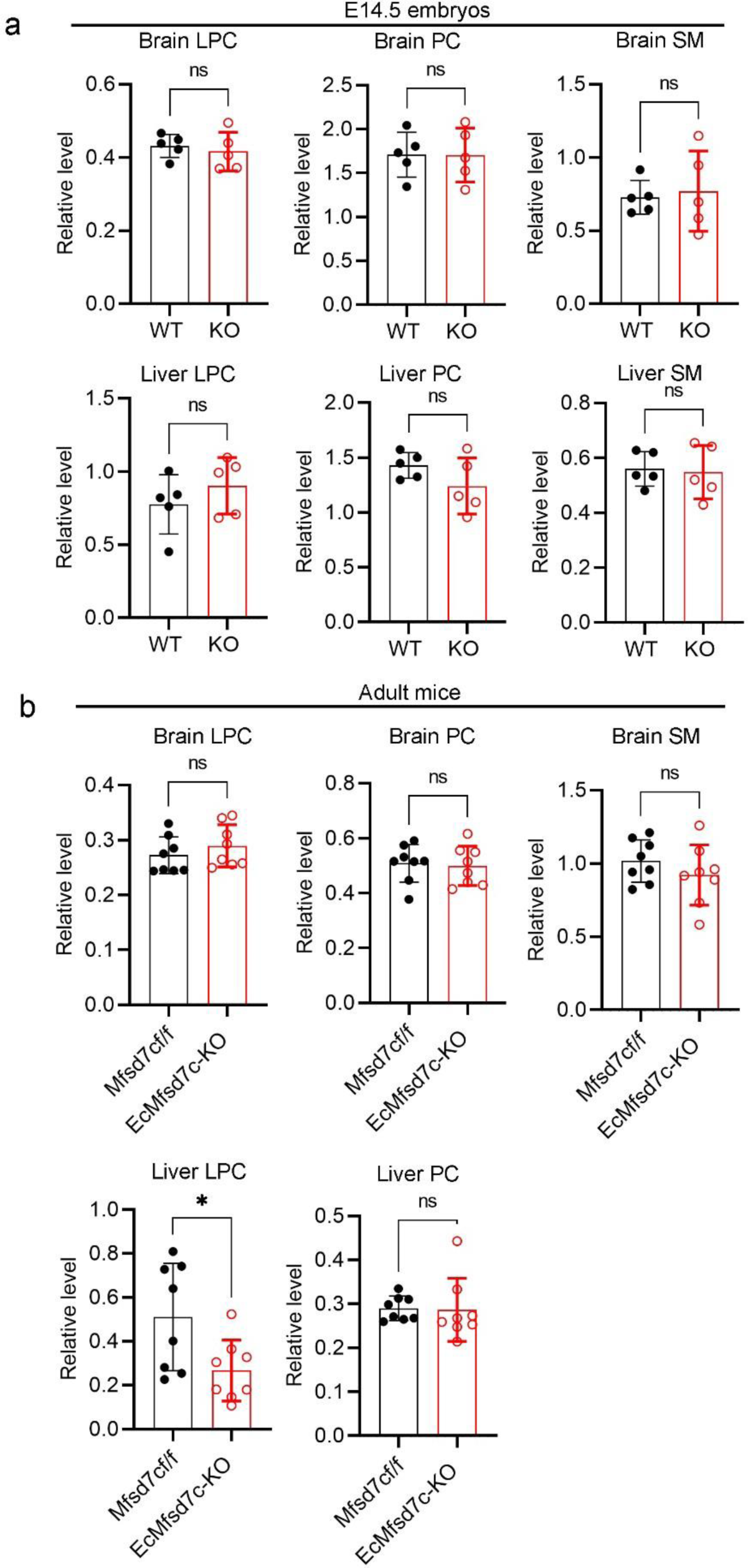
Deletion of Mfsd7c did not affect the levels of endogenous phospholipids in the brain and liver. **a,** Lipidomic analysis of phospholipids containing choline including LPC, PC, and SM in the brains and livers of E14.5 Mfsd7c KO and wild-type littermates. **b,** Lipidomic analysis of LPC, PC, and SM in the brains and livers from adult EcMfsd7c-KO and control mice. Each symbol represents one mouse. Data are expressed as mean ± SD. *P<0.05; t-test was used. Full list of phospholipids can be found in the **supplemental data tables 13-16.**

**Extended Data Fig. 8.**
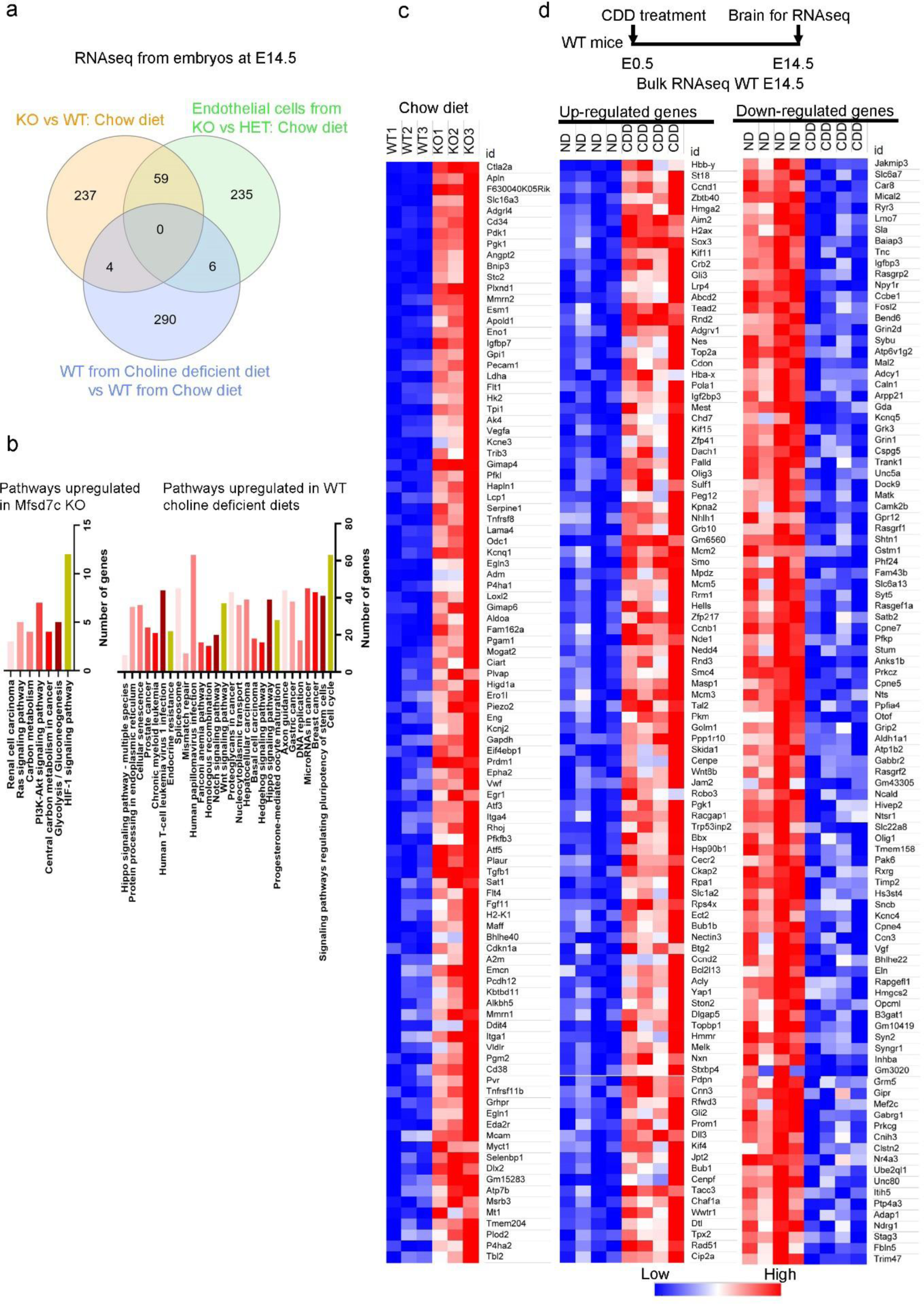
Gene expression changes due to maternal choline deficiency did not recapitulate the loss of Mfsd7c in the embryos. **a,** Venn diagram of differential gene expression change in the brains of Mfsd7c knockout embryos and wildtype embryos from maternal choline deficiency. In comparison amongst the top 300 genes that were differentially changes in the whole brain and isolated CNS endothelial cells from Mfsd7c knockout, there were a significant overlap. However, there was a few genes that were changed due to choline deficiency compared to the lack of Mfsd7c. **b,** Upregulated biological pathways in the brain of Mfsd7c KO embryos from mothers fed with choline sufficient diet and wild-type embryos from mothers fed with choline deficient diet. **c,** Heatmaps of upregulated genes from Mfsd7c KO compared to wild-type littermates. **d,** Heatmaps of upregulated and downregulated genes from wild-type embryos from mothers fed with choline deficient diet compared to wild-type embryos from mothers fed with choline sufficient diet. There is no major overlap of upregulated pathways due to Mfsd7c deletion and choline deficiency. The RNAseq dataset for WT from ND and CDD can be found in the supplemental data tables 17.

**Extended Data Fig. 9.**
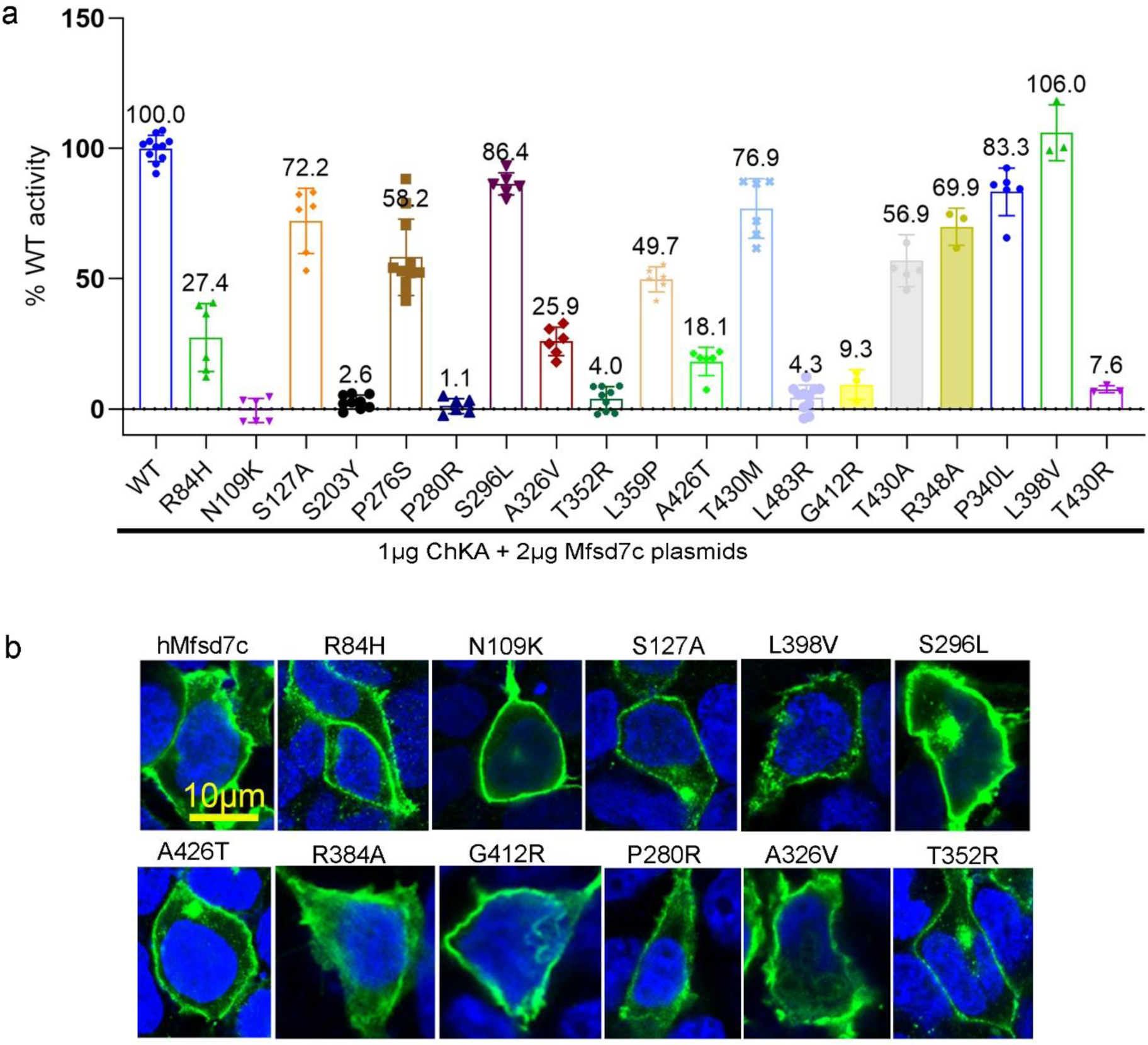
Missense mutations of Mfsd7c affect choline transport activity. **a,** Transport activity of the missense mutations of Mfsd7c that were reported previously in the literature. Transport activity of these missense mutations was expressed as percentage of wildtype (WT) protein. Each symbol represents one replicate. Experiments were repeated at least twice. **b**, Fluorescent microscope analysis of the localization of the missense mutations in HEK293 cells. Localization of these missense mutations to the plasma membrane as similar to the wild-type protein was not affected. A full list of mutants can be found in the **supplemental data table 18.**

## Notes

### Competing Interest Statement

The authors have declared no competing interest.

### Summary of Updates

Figure 5f & g

